# Single-cell transcriptomic atlas of frontoinsular cortex reveals molecular correlates of selective neuronal vulnerability in FTD

**DOI:** 10.64898/2026.07.13.738307

**Authors:** Arnar Breevoort, Dimitar Ivanov, Liam Horan-Portelance, Alissa Nana, Sarat Vatsavayai, Àlex Tudoras Miravet, Jenelle L. Wallace, Jingwen W. Ding, Felipe L. Pereira, Kristen Fernhoff, Maria Luisa Gorno-Tempini, Salvatore Spina, Jennifer S. Yokoyama, Howard J. Rosen, Lea T. Grinberg, Frank M. J. Jacobs, Bruce L. Miller, William W. Seeley, Alex A. Pollen

**Author notes:** These authors contributed equally: Liam Horan-Portelance, Dimitar Ivanov.

## Abstract

Frontotemporal dementia (FTD) is characterized by selective neuronal vulnerability, yet the features that predispose specific neuron types to degeneration remain unclear. We performed single-nucleus RNA sequencing of frontoinsular cortex, a region affected early in behavioral variant FTD, across individuals with *C9orf72*-associated and sporadic FTD-MND spectrum disease. By enriching for large projection neurons, we resolved molecular subtypes of layer 5 extratelencephalic neurons, including von Economo neurons, and identified selective depletion of specific layer 2/3 and layer 5 neuron subtypes, convergent across genotypes. Despite selective neuronal loss, disease-associated transcriptional changes were convergent across excitatory neuron populations, suggesting that they reflect upstream pathophysiology or shared responses to local neurodegeneration. By relating neighborhood-level depletion in disease to gene expression in controls, we found that baseline cellular respiration and ATP synthesis predict neuronal vulnerability in disease. These findings define molecular correlates of selective neuronal vulnerability in FTD and provide a framework linking cell type and state to neurodegeneration.

## Introduction

Behavioral variant frontotemporal dementia (bvFTD) and motor neuron disease (MND) are progressive neurodegenerative disorders linked by a shared TDP-43 proteinopathy and genetic risk factors^1,2^. Despite their close molecular-genetic ties, regional pathology and neuronal loss begin in a syndrome-specific manner, with frontal paralimbic atrophy in bvFTD^3–9^ and pyramidal motor system degeneration in MND^10,11^. Within these systems, however, both disorders are linked to early degeneration of large, morphologically distinct and region-specific layer 5 projection neurons: von Economo neurons (VENs) and fork cells of the anterior cingulate and frontoinsular cortices in bvFTD and precentral gyrus upper motor neurons, including Betz cells, in MND^4,5,12^. These observations suggest that selective vulnerability may reflect shared intrinsic properties of specialized projection neuron subtypes, but the molecular basis of this vulnerability remains unknown.

Recent single-nucleus transcriptomic studies have begun to characterize cellular responses to neurodegeneration, identifying shared and cell type-specific gene expression changes across brain regions and pathological entities. In the context of FTD/MND, however, these studies have 1) largely focused on prefrontal and primary motor cortices^6,13,14^, 2) undersampled the most vulnerable neuron types, and 3) defined vulnerability by the magnitude of transcriptional changes observed in disease. As a result, the precise molecular identities of the most depleted neuronal subtypes and the underlying transcriptional programs that predispose to vulnerability remain unclear, especially for bvFTD. Despite these caveats, one study showed that Layer 5 extratelencephalic (L5 ET) neurons, the class that includes VENs, fork cells, and Betz cells, are the most transcriptionally abnormal of any cell type brain-wide^6^, and another showed that L5 ETs are the most predisposed cell type to cryptic splicing events caused by loss of TDP-43 function^15^.

The frontoinsular cortex (FI) plays vital roles in social, emotional, and autonomic/interoceptive functions^16–18,19,20,21^. FI is among the earliest sites of neurodegeneration in bvFTD^3–9,22,23^ and contains the highest density of von Economo neurons (VENs) and fork cells^19^, yet no single-cell map of FI exists for bvFTD or MND patients. Here, we generated a single-nucleus transcriptomic atlas of FI across 40 individuals including controls and patients with *C9orf72*-associated and sporadic FTD/MND. By enriching for large neurons, we achieved the largest single-nucleus census of human L5 ET neurons to date, which we used to resolve previously unrecognized molecular subtypes. We find convergent transcriptional remodeling across excitatory neuron populations in disease, alongside selective depletion of specific subtypes of L2/3 and L5 neurons. Importantly, we identify gene expression programs present in control neurons—including metabolic and synapse assembly pathways—that predict neuronal loss in disease. These findings define intrinsic features of selectively vulnerable neurons and provide a framework for understanding cell type-specific degeneration in FTD/MND.

## Results

### Molecular cell type diversity in frontoinsular cortex

Although recent studies explored the transcriptional consequences of FTD/MND in the middle frontal and precentral gyri^6,14^, here we focused on FI, the most consistently and often the earliest affected brain region in patients with bvFTD^22,24^ and the only region studied to date that contains more than a scattering of VENs and fork cells. We analyzed frozen post-mortem FI collected from 40 participants by the UCSF Neurodegenerative Disease Brain Bank. To reduce pathological and genetic heterogeneity, we limited enrollment to patients falling along the clinical bvFTD-MND spectrum who had *C9orf72*-associated (N = 16) or sporadic (N = 15) FTLD/MND-TDP Type B (or an unclassifiably sparse subtype, “Type U”, in some patients) and controls (N = 9). Within this cohort, patients showed a range of FI-relevant clinico-pathological involvement from few if any bvFTD symptoms and no TDP-43 inclusions outside the motor system (in MND cases) to advanced bvFTD with severe FI degeneration and extensive TDP-43 pathological burden (Fig. 1a, Suppl. Table 1). Patients with MND and those with FTLD-TDP Type U were included in hopes of capturing the earliest stages of TDP-43-related FI degeneration. To mitigate batch effects, we selected samples with comparable post-mortem intervals (mean PMI: 10.7h vs 14.3h; SD 6 vs 7.4) and processed tissue samples together in a common pool for nuclei isolation (Fig. 1b).

**Fig. 1.**
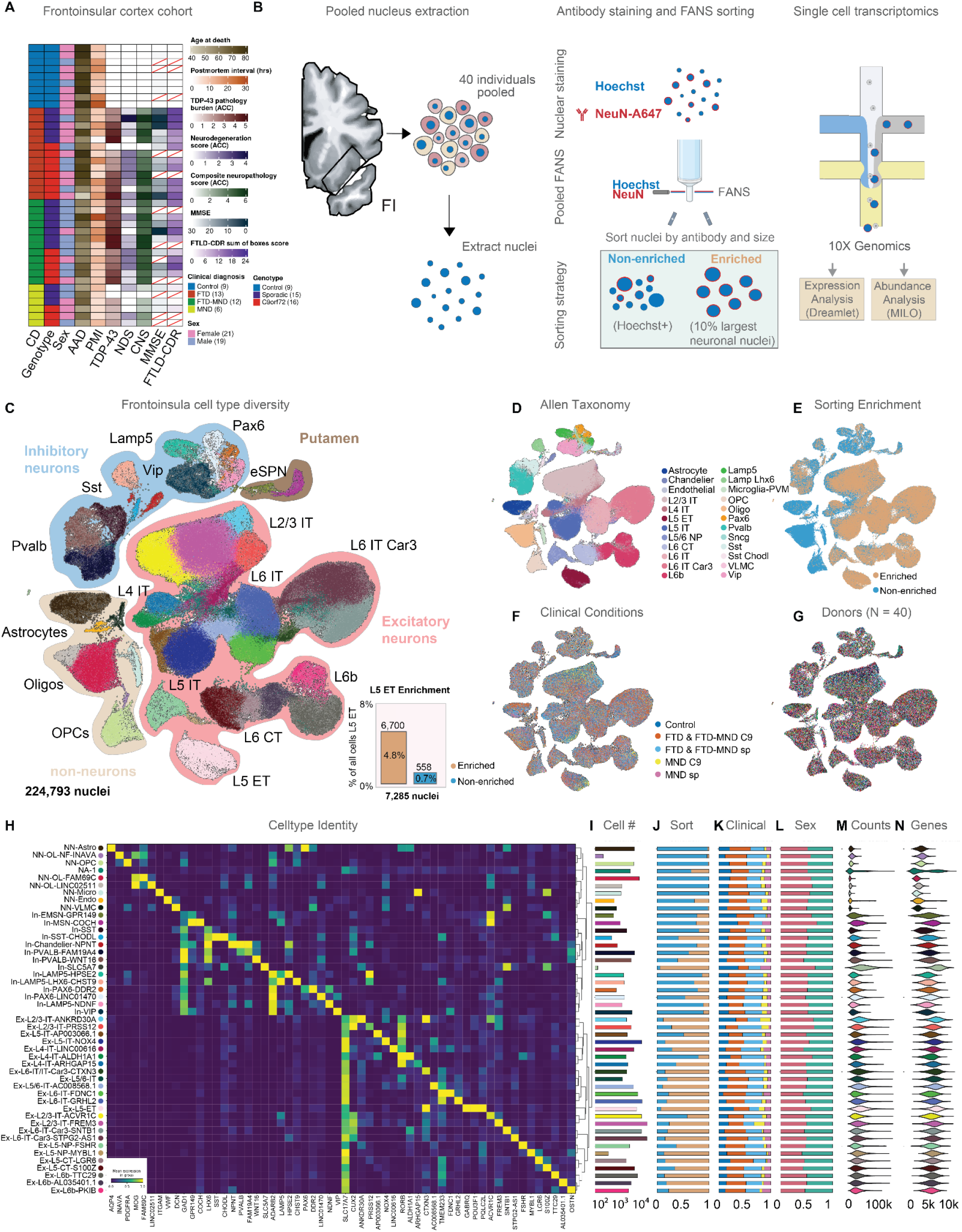
Molecular cell type diversity of frontoinsular cortex in pathologically normal and bvFTD and MND patients. **A:** Clinical, demographic, and neuropathological characteristics of the frontoinsular cortex (FI) cohort comprising 40 postmortem samples from controls and patients spanning the bvFTD-MND spectrum, including *C9orf72*-associated and sporadic FTLD/MND-TDP. Full donor metadata, including a suite of clinical and pathological measures, can be found in Suppl. Table 1a. Missing data is indicated with a red slash. **B:** All 40 samples were pooled to reduce batch effects. Nuclei were FANS-sorted into a non-enriched Hoechst+ condition and a size-gated enriched condition comprising the largest 10% of neuronal nuclei by NeuN signal and FSC-A gating (“LN10”), then processed using 10x Genomics 3’ single-cell sequencing. Differential expression and differential abundance analyses were performed using Dreamlet and Milo, respectively. **C:** UMAP visualization of the integrated FI snRNA-seq dataset showing major neuronal and non-neuronal populations across the full spectrum of FI cellular diversity (47 cell populations; cell level 3 taxonomy). The LN10 enrichment strategy substantially increased recovery of L5 ET neurons (n = 7,285) relative to un-enriched. Inset shows enrichment of L5 ET neurons in the LN10 sort. A total of 224,793 nuclei passed quality control. **D:** Reference-based annotation mapping of FI cell populations using Allen Brain Cell Atlas taxonomies (Suppl. Table 1b; cell_level_2). **E:** UMAP colored by sorting condition showing enrichment of excitatory neuron populations in the LN10 sort. **F-G:** UMAPs colored by clinical condition (F) and donor identity (G), demonstrating robust integration across disease groups and individuals. **H:** Hierarchical clustering and marker gene expression across the 47 transcriptionally defined FI cell populations (cell level 3 taxonomy). **I-J:** Cell numbers and relative proportions from the Hoechst+ and LN10 sorting conditions across FI cell populations. **K:** Relative proportions of cells from each clinical disease group across FI cell populations: Control (blue), *C9orf72* FTD & FTD-MND (orange), sporadic FTD & FTD-MND (cyan), *C9orf72* MND (yellow), and sporadic MND (pink). **L:** Relative proportions of cells from female and male donors across FI cell populations. **M-N:** Transcript and detected gene distributions across FI cell populations. Neuronal populations exhibited higher transcript and gene counts relative to non-neuronal populations.

Recent transcriptomics studies provide unbiased support for selective L5 ET vulnerability but undersampled this rare neuronal population, limiting insights into factors that underlie its selective degeneration^6,25^. Enrichment strategies targeting selectively vulnerable neuronal populations have increased resolution of relevant cell types in Alzheimer’s^26^, Parkinson’s^27^ and Huntington’s disease^28^, but have not been applied to FTD/MND. To address this barrier, we enriched for the largest ∼10% of neurons (LN10) using fluorescence activated nucleus sorting (FANS) alongside an unenriched Hoechst+ condition (Fig. 1b, Extended Data Fig. 1b-g)^29^. Comparing LN10 enrichment to Hoechst across neighborhoods of transcriptionally similar cells, defined using the Milo framework^30^, provided a proxy for nucleus size based on differential abundance across the Hoechst+ and LN10 conditions (Extended Data Fig. 1h). We performed single nucleus RNA sequencing (snRNA-seq) using 10x Genomics V3.1 HT kits and Illumina sequencing, recovering 224,793 total cells after quality control (85,635 Hoechst+; 139,158 LN10; see Methods). To assign cells to individual-of-origin and estimate ambient RNA contamination, we leveraged naturally occurring genetic variation using CellBouncer^31^, recovering all individuals (mean cells per individual: 5,620, SD: 3,126) with limited ambient RNA across lanes (mean ambient RNA score: 0.07, SD: 0.05).

To analyze the molecular diversity of FI cell types, we integrated data across individuals using single cell variational inference (scVI)^32^ (Methods). As expected, we observed excitatory neurons, inhibitory neurons, and non-neural cells (Level 1 annotations) (Fig. 1c). Reference mapping via MapMyCells (RRID:SCR_024672) to the Allen 10x Whole Human Brain taxonomy and the 10x Human MTG SEA-AD taxonomy^25,26^ provided general class labels for interpreting FI cell type diversity, with the LN10 condition enriching for excitatory neurons (Level 2 annotations; Fig. 1e). We next performed Leiden clustering, retaining 47 clusters with qualitatively distinct markers, with cells from different individuals and clinical, pathological, and genetic subtypes distributed across clusters (Level 3 annotations; Fig. 1c-g, Extended Data Fig. 2a-c, Methods). Among non-neuronal cells, we observed clusters representing different stages of oligodendrocyte lineage differentiation, astrocytes, microglia, and vascular cells. Among inhibitory neurons, we observed 14 clusters, corresponding to cardinal classes of cortical inhibitory neurons, including SST, PVALB, VIP and LAMP5 subtypes^33^. In addition, we observed two clusters of LGE-derived medium spiny projection neurons (In EMSN GPR149 & In MSN COCH) and a small cluster of cholinergic interneurons (In SLC5A7), likely from adjacent putamen (Extended Data Fig. 2a).

Excitatory neuron subtypes vary across cortical areas^34^, but we lack a detailed account of excitatory neuron diversity in human FI. Our enrichment strategy for large neuronal nuclei provided additional excitatory neuron subtype resolution, including a 7.5-fold enrichment for L5 ET neurons (Fig. 1c, Extended Data Fig. 2a). We observed four clusters mapping to L2/3 intratelencephalic (IT) excitatory neurons and three clusters mapping to L4 IT neurons. As FI is cytoarchitecturally agranular, these L4 IT cells likely fall in layers 3 and/or 5 or a histologically indistinct L4 as observed in M1^35^, but we retained the L4 IT label for taxonomic consistency. We further observed diverse L5 and L6 subtypes, including nine presumptive IT subtypes, two corticothalamic (CT) subtypes, two near projecting (NP) subtypes, and three L6b subtypes. Based on the expression of *CPLX3*, *CABP7*, and *CTGF*, the L6b subtypes correspond to recently observed populations in proposed to be homologous to the claustrum shell^36^, while the three populations mapping to L6 IT Car3 cells may include claustrum populations based on close homologies observed in mouse and proximity of claustrum to FI^37^.

The size-based enrichment had the most striking effects on the abundance of excitatory neurons in L2/3. To interpret the increased diversity among L2/3 cell types in our dataset, we mapped our transcriptomic data to a multimodal catalogue of middle temporal gyrus (MTG), where cell size, position, morphology and physiology were recently cataloged across MTG populations^38^. The L2/3 ACVR1C population was strongly depleted among large nuclei in our dataset (Extended Data Fig. 3a-b) and best corresponded to the L2 LAMP5 LTK population in MTG (Extended Data Fig. 3c,d,f), consistent with the small size and superficial location of this homologous population in MTG (Extended Data Fig. 3e). In contrast, among excitatory L2/3 IT FREM3 cells, we observed a gradient of LN10 enrichment (Extended Data Fig. 3b) consistent with the size and depth gradient of Ex L2/3 FREM3 cells described in MTG, in which larger cells with complex dendritic arbors were reported deeper in L3. Similarly, the L2/3 IT ANKRD30A population, although lacking a clear homolog in MTG (Extended Data Fig. 3d,f), shows high expression of *NEFL* and *NEFH* (Extended Data Fig. 3g), neurofilament genes that encode proteins labeled by SMI-32 in large, long-range projecting layer 3 neurons^38,39^. This population is additionally distinguished by selective expression of *SYT2* (Extended Data Fig. 3g), a synaptic vesicle calcium sensor associated with rapid neurotransmitter release, also expressed in L5 ET neurons and other long-range projecting cell types. Together, these data provide a high-resolution map of FI cellular diversity integrated across pathologically normal and disease samples with enhanced representation of large projection neurons, providing a reference for examining the cell type-specific transcriptional consequences and cellular vulnerabilities in FTLD/MND-TDP with molecular subtype resolution.

### Convergent upregulation of dendritic, synaptic and axonal gene programs in *C9orf72* and sporadic bvFTD

We next examined the landscape of transcriptional changes seen in *C9orf72* and sporadic bvFTD/MND across molecularly defined cell types. In line with recent reports^6^, we hypothesized that *C9orf72* and sporadic bvFTD would display convergent transcriptional consequences reflecting shared disease processes. Additionally, given the paucity of FI pathological changes in clinically pure MND, we predicted that transcriptional changes in MND-only participants would represent incipient FTD, considering that FI TDP-43 pathobiology is often just beginning to emerge in patients with pure MND. To mitigate the issues of pseudoreplication and inherent sparsity of transcript detection at the level of individual cells, we applied precision-weighted linear mixed models using Dreamlet^40^ to pseudobulk expression profiles aggregated by individual, tested across clinical subtype and genotype, and controlled for sex, age at death, and postmortem interval (PMI) (Methods). This model was applied to both our most coarse (Level 1) and most granular (Level 3) cellular taxonomy (Extended Data Fig. 4a-g & Suppl. Table 1b).

We first compared the broad landscape of differential gene expression across four major disease conditions using the course cellular taxonomy (Level 1). We grouped clinical bvFTD and bvFTD-MND together given their shared involvement of FI and split them into *C9orf72* and sporadic groups; MND was treated separately, also divided by genotype. We calculated Pearson correlations of log_2_ fold changes (logFC in cases versus controls) across all broad cell types tested (Level 1) for all genes (Fig. 2a). We observed positive correlations between all conditions, with the strongest concordance between *C9orf72* and sporadic bvFTD & bvFTD-MND, as expected (Pearson’s r = 0.79) (Fig. 2a). Pure MND cases showed weaker, albeit positive, correlations with both bvFTD/bvFTD-MND cases and each other, reflecting the smaller absolute effect sizes in pure MND cases (Fig. 2a). Given that pure MND showed weaker effects that correlated less well with the bvFTD groups, most downstream analyses included only cases with clinical bvFTD (i.e., bvFTD or bvFTD-MND, henceforth “FTD”).

**Fig. 2.**
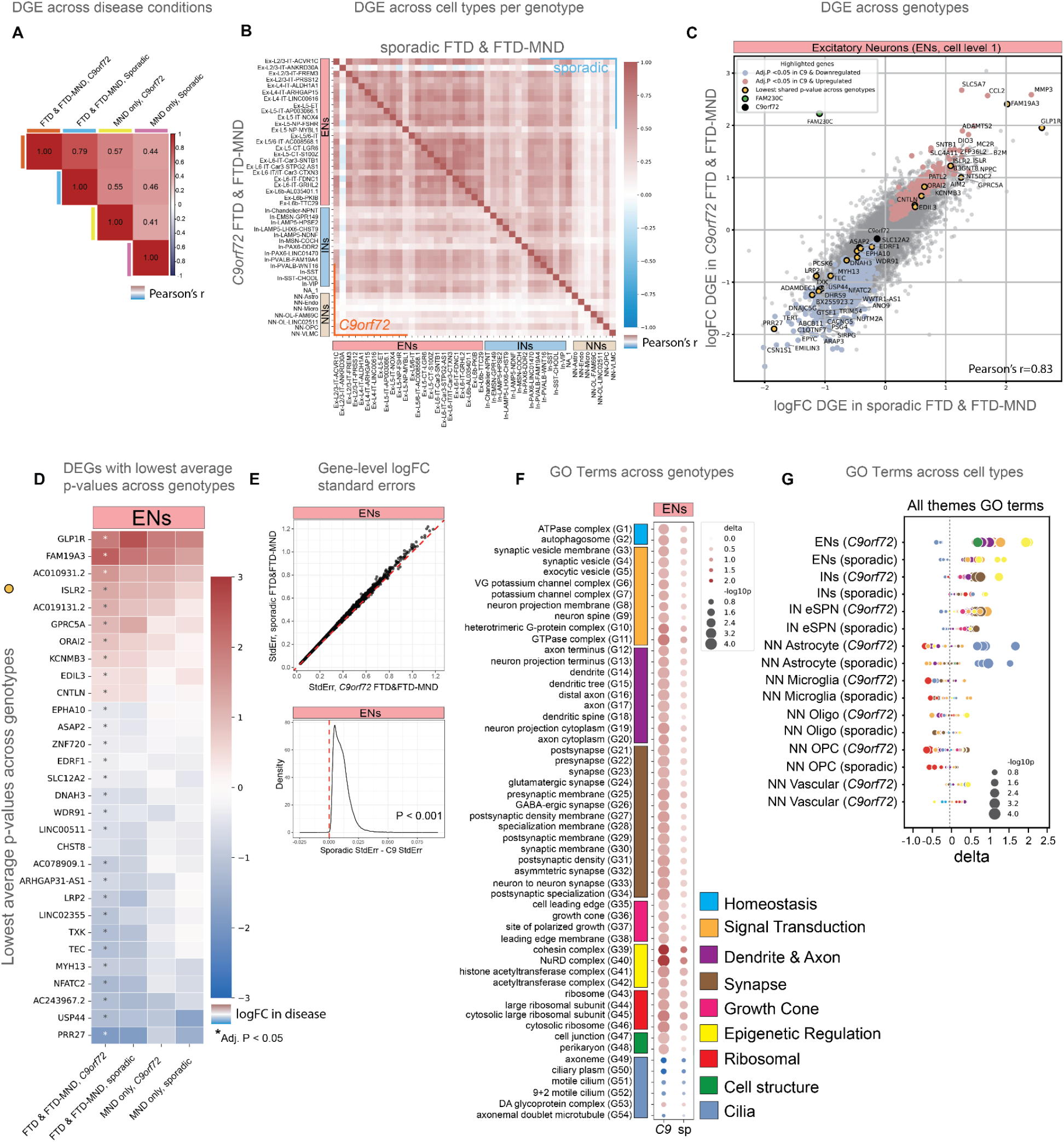
Convergent upregulation of dendritic, synaptic and axonal gene programs in *C9orf72* and sporadic FTD & FTD-MND. **A:** Disease-associated differential gene expression (DGE) logFC values were positively correlated across clinical and genetic disease groups. *C9orf72* FTD & FTD-MND shown in orange, sporadic FTD & FTD-MND shown in blue, *C9orf72* MND shown in yellow, and sporadic MND shown in pink. **B:** Pearson correlation matrix of disease-associated DGE logFC values across transcriptionally defined FI cell populations within *C9orf72* and sporadic FTD & FTD-MND separately. Correlations among sporadic FTD & FTD-MND cell populations are shown in the upper-right triangle, whereas correlations among C9orf72 FTD & FTD-MND populations are shown in the lower-left triangle. Only genes which passed filtering criteria across all assays were used to calculate pairwise correlations. Broadly shared transcriptional responses were observed across cell populations, with the strongest convergence among excitatory neuron (EN) subclasses and more modest convergence among inhibitory neurons (INs) and non-neuronal populations. **C:** Comparison of disease-associated DGE logFC values in ENs (level 1) between *C9orf72* and sporadic FTD & FTD-MND. Disease-associated transcriptional changes were highly concordant across genotypes (Pearson’s r = 0.83; P < 0.05). Highlighted genes in blue or salmon met significance thresholds in one or both disease groups, and genes in yellow are coding genes from the top 30 DEGs with the lowest average adjusted P values across genotypes. *FAM230C* (highlighted in green) was the only gene significantly differentially expressed when directly comparing *C9orf72* and sporadic FTD & FTD-MND (sporadic vs. *C9orf72*, logFC = -3.31, adj. P = 0.0094). **D:** LogFC values of the 30 most significantly differentially expressed genes in FTD & FTD-MND, selected by lowest average adj. P across *C9orf72* and sporadic FTD & FTD-MND. The direction of differential expression was generally shared across FTD and MND disease groups. **E:** Comparison of gene-level standard errors of disease-associated logFC estimates in ENs between *C9orf72* and sporadic FTD & FTD-MND. Sporadic cases exhibited significantly higher variance in DGE estimates relative to *C9orf72* cases (P < 0.001). **F:** Gene ontology enrichment analysis across genotypes in ENs showing convergent upregulation of synaptic, dendritic, axonal, growth cone, and epigenetic programs, alongside downregulation of cilia-related pathways in FTD & FTD-MND. **G:** Gene ontology enrichment analysis across broad FI cell classes (Suppl. Table 1b; cell_level_1). Neuronal structural and synaptic pathway upregulation was most prominent in ENs, whereas cilia-related pathways were broadly downregulated across most cell populations but uniquely upregulated in astrocytes.

We next performed the same analysis, instead comparing gene expression across the most granular cell type annotations (Level 3) within *C9orf72* and sporadic FTD cases separately (Fig. 2b). This revealed broadly correlated gene expression changes across cell types within each genotype, particularly evident in excitatory neuron (EN) subtypes (Fig. 2b). Given these shared patterns among excitatory neuron classes, we grouped ENs to increase statistical power, observing strongly correlated changes in magnitude and directions of effects between genotypes (Pearson’s r = 0.83) (Fig. 2c). Intriguingly, when evaluating most significant DEGs observed in FTD, the pure MND cases showed directionally similar effects, albeit with smaller effect sizes that did not pass our stringent statistical thresholds (Fig. 2a,d), likely due to sample size and milder FI involvement. Despite correlated expression changes and comparable cell numbers, many more significant DEGs (adj.p<0.05) were observed in *C9orf72* compared to sporadic FTD cases, with 818 vs. 0 DEGs in ENs at level 1, and 2,981 vs. 293 DEGs across all 47 cell types at level 3, in *C9orf72* vs. sporadic FTD, respectively (Suppl. Table 2a,b). Analysis of gene-level standard errors of disease-related log_2_ fold changes (logFC) in the FTD cohort revealed higher variance among sporadic cases than *C9orf72* cases (Fig. 2e), perhaps reflecting the more heterogeneous and still unknown origins of sporadic FTLD/MND-TDP. Shared correlations between case versus control logFC across genotypes were also found in inhibitory neurons and non-neuronal cell types such as astrocytes, oligodendrocytes, OPCs, microglia and vascular cells (Extended Data Fig. 5a-f), pointing towards convergence of disease-related gene expression patterns across both genotypes and cell types. Among all cell types, we found the most DEGs, in decreasing order, in ENs, astrocytes, oligodendrocytes, and INs (Extended Data Fig. 5g-h & Suppl. Table 2).

In excitatory neurons, upregulated genes included factors involved in axonal growth, synaptic function, cellular adhesion, such as *ISLR2*^41^, *CNTLN*^42,43^, and *EDIL3*, consistent with remodeling of neuronal structure and connectivity. Notably, *GLP1R,* a receptor regulating satiety and the target of appetite-curbing GLP1R agonist medications^44^, was upregulated across genotypes, driven by an expansion in the proportion of excitatory neurons expressing *GLP1R*, with the strongest effects observed in L2/3 IT, L5 ET and L5/6 IT populations (Fig. 2c; Extended Data Fig. 6a). Altered satiety cues may drive overeating behaviors that represent a core feature of bvFTD^45^. Additional highly upregulated genes included *MMP3*, *ADAMTS2*, *CCL2*, *FAM19A3* and *SLC5A7*, reflecting extracellular matrix remodeling, inflammatory signaling, and cholinergic pathway engagement (Fig. 2c).

Downregulated genes included regulators of synaptic organization and neuronal maintenance, such as *EPHA10*, *LRP2*^46^, and *ARAP3*^47^, alongside genes linked to neuroinflammation, including *TXK*^48^. The most strongly downregulated gene, *CSN1S1,* was previously identified as a precentral gyrus L5 ET/Betz cell marker^49^. Additionally, deubiquitinating enzyme gene *USP44*^50^, which regulates neurite outgrowth *ARAP3*^47^, and telomerase reverse transcriptase *TERT,* which prevents telomere erosion and maintains levels of anti-oxidative enzymes^51^ were also strongly downregulated. Although the overall direction of differential expression was largely shared across genotypes, FAM230C showed significant discordant effects between genotypes (sporadic vs. *C9orf72*, logFC = -3.31, adj. P = 0.009) (Fig. 2c).

To further evaluate pathways disrupted in FTD, we performed gene ontology enrichment analysis. Consistent with a recent study in middle frontal and precentral gyri^6^, EN cilia-related gene programs were downregulated in FTD (Fig. 2f). Against a background of downregulation in most cell types (ENs, INs, microglia, oligodendrocytes, OPCs and vascular cells), however, we observed strong cilia program upregulation in astrocytes (Fig. 2g & Extended Data Fig. 7a,c). Genes driving these cilia programs included *DNAI1*, *CFAP53*, and *IQUB* (Extended Data Fig. 5i). Again, cilia gene modulation was correlated across genotypes but reached higher significance levels in *C9orf72*.

The most pervasive pathway alteration in FTD reflected convergent upregulation of dendritic, axonal, synaptic, and epigenetic regulation gene programs in ENs (Fig. 2f-g & Suppl. Table 2c). This upregulation was also broadly shared across individual EN subtypes, and to a lesser extent INs (Extended Data Fig. 7c). Genes driving these enrichments, as defined by passing significance in the Dreamlet analysis (Level 1), included the choline transporter, *SLC5A7*, nerve growth factor receptor, *NGFR*, the opioid receptor, *OPRK1*, and the cell adhesion gene *EFNB1* (Extended Data Fig. 5i). Whether these findings reflect a compensatory response or a pathological process remains to be determined.

Together, these findings indicate a widespread and coordinated transcriptional remodeling of neuronal structural and synaptic programs in FTD that occurs broadly across neuronal populations. Consistent with prior reports of pan-neuronal upregulation of axonal damage response genes in ALS and FTLD^6^, and synaptic program upregulation in upper-layer neurons in ALS^52^, our findings extend this observation to a broader set of structural, synaptic, and epigenetic programs in FTD.

### Convergent L2/3 IT, L5 IT, and L5 ET depletion in *C9orf72* and sporadic bvFTD

Given the highly convergent transcriptional consequences and shared pathological features of *C9orf72* and sporadic FTD/MND, we next examined the extent of shared and specific cell type vulnerabilities. Classical histology studies indicate that VENs, fork cells, possibly other L5 ETs, and, to a lesser extent, L2/3 IT neurons, exhibit increased propensity for TDP-43 aggregation and death in bvFTD^3–97,8,23,53^. A recent unbiased, single nucleus transcriptomic study supported L5 ET vulnerability in FTLD-TDP via measures of transcriptional dysregulation^6,54^. However, molecularly defined subpopulations showing depletion, and the baseline transcriptomic signatures that predict vulnerability across and within such populations, remain to be elucidated. To examine cellular vulnerabilities agnostic to cluster boundary definitions, we performed differential abundance testing across transcriptionally similar cell neighborhoods, controlling for remaining variation in clinical diagnosis, genotype, sex, age at death, and PMI using the Milo statistical framework^30^ (Methods). This neighborhood-based approach enables a more granular view of vulnerability than cluster-level analyses and provides a continuous representation of cell states across which disease-related depletion can be assessed.

Consistent with the neuropathological literature, we observed marked depletion among L2/3 IT and L5 ET neurons in *C9orf72* and sporadic cases (Fig. 3a-b). Interestingly, depleted neighborhoods were distributed unevenly within these broad classes. Within L2/3 IT neurons, depletion was concentrated in specific subpopulations, including L2/3 IT ANKRD30A and a subset of L2/3 IT FREM3, whereas L2/3 IT ACVR1C neighborhoods were relatively spared (Fig. 3a–b; Suppl. Table 3a). In addition to strong depletion of L5 ET neurons, we observed heterogeneous changes in abundance among L5 IT NOX4 neighborhoods, with a subset of neighborhoods showing pronounced depletion while others remained relatively resistant (Fig. 3a-b). With the exceptions noted above, other deep layer neuron types, regardless of projection target (i.e., L5/6 IT, L6 IT, L6 IT CAR3, L6 CT, and L6b), were for the most part resistant to cell loss (Fig. 3a-b).

**Fig. 3.**
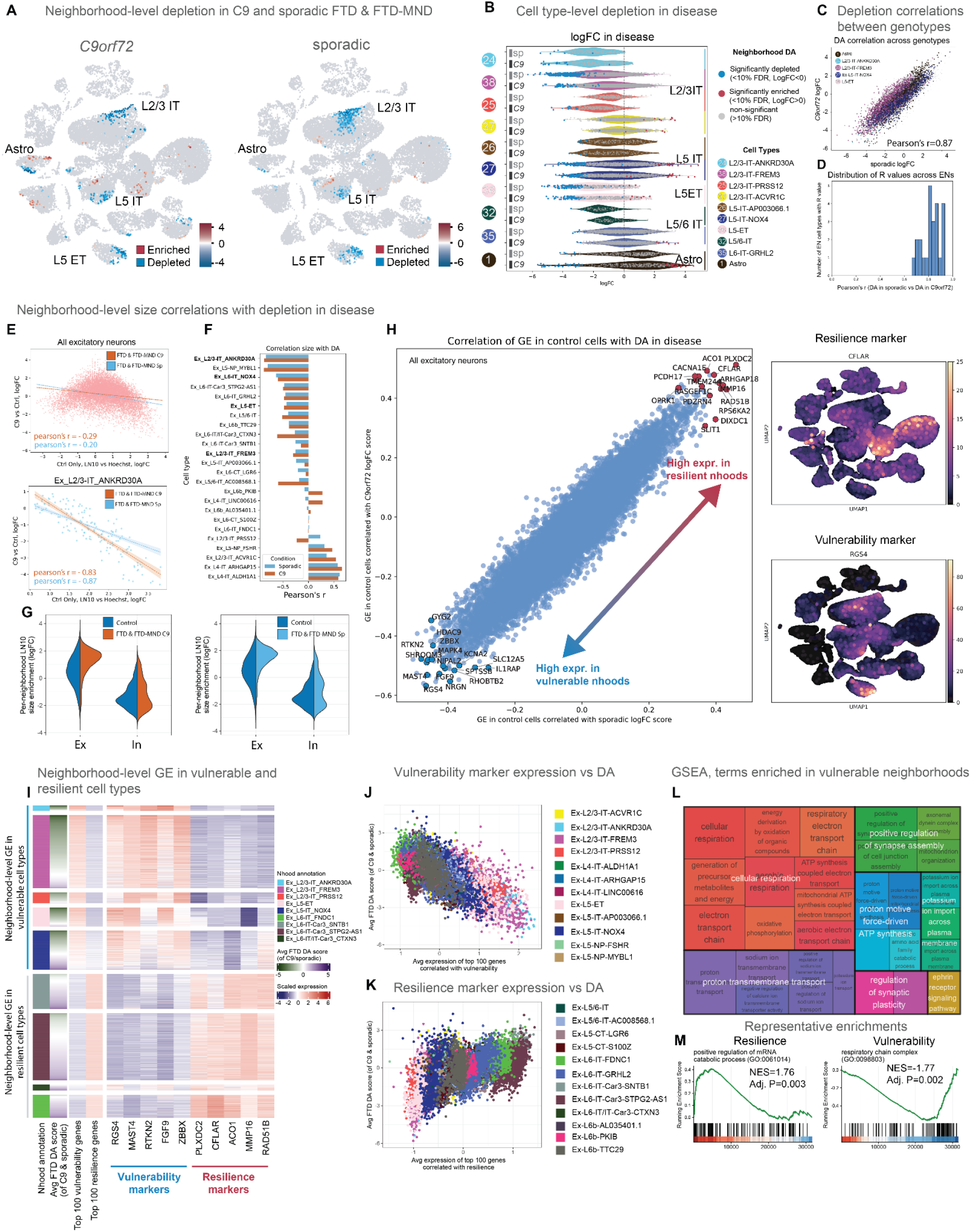
Convergent L2/3 IT. L5 IT, and L5 ET depletion in *C9orf72* and sporadic bvFTD. **A:** Neighborhood-level differential abundance (DA) from *Milo* analysis in *C9orf72* and sporadic FTD & FTD-MND is convergent across genotypes, and depletions (blue) or enrichments (red) in disease are most prominent in L2/3 IT, L5 IT, L5 ET and Astrocyte classes. Only neighborhoods with significant depletion or enrichment in disease (FDR < 0.1) are highlighted. **B:** Distribution of neighborhood-level DA logFC values across FI cell populations (level 3) in *C9orf72* and sporadic FTD & FTD-MND. Significant depletion was concentrated within L2/3 IT ANKRD30A, L2/3 IT FREM3, L5 IT NOX4, and L5 ET populations, whereas other excitatory neuron populations remained relatively resistant. **C**: Neighborhood-level DA values were highly concordant across *C9orf72* and sporadic FTD & FTD-MND. Shown are neighborhood-level DA values for the most affected cell types from panel B (Pearson’s r = 0.87). **D:** Distribution of Pearson correlation coefficients comparing neighborhood-level DA values between *C9orf72* and sporadic FTD & FTD-MND across excitatory neuron populations (Suppl. Table 1b; cell_level_3), demonstrating strong concordance of vulnerability patterns across genotypes. **E:** Relationship between LN10 enrichment score, used as a proxy for neuronal size, and disease-associated DA across excitatory neuron neighborhoods reveals modest correlation of nuclear size with disease-associated DA across all ENs (Pearson’s r = -0.29 and -0.20 for *C9orf72* and sporadic FTD & FTD-MND, respectively), indicating nuclear size does not strongly explain EN vulnerability across all cell types. However, a strong correlation exists for L2/3 IT ANKDR30A neighborhoods (Pearson’s r = -0.83 and -0.87), indicating nuclear size largely explains their vulnerability. Note that for both sub-panels, only *C9orf72* points are plotted, but trend lines for both genotypes are overlaid. **F:** Correlation coefficients between neighborhood-level LN10 enrichment and disease-associated DA across excitatory neuron subclasses in *C9orf72* and sporadic FTD & FTD-MND. Most vulnerable excitatory neuron populations from panel B are highlighted in bold. **G:** Distribution of large nuclei (LN10 fraction) neighborhood-level enrichment scores in patients and controls for ENs and INs. Excitatory neurons exhibited increased LN10 enrichment in disease, indicative of increased nuclear size, whereas inhibitory neurons showed comparatively limited changes. **H:** Correlation between baseline gene expression in control excitatory neuron neighborhoods and disease-associated DA in *C9orf72* and sporadic FTD & FTD-MND identifies vulnerability-associated (blue) and resilience-associated (red) genes. Feature plots show the strongest resilience-associated (*CFLAR*) and vulnerability-associated (*RGS4*) genes. **I:** Average expression of the 100 most strongly correlated vulnerability and resilience genes across neighborhoods shows high vulnerability marker expression in depleted cell types and high resilience marker expression in enriched cell types. Plotted are DA logFC values in disease per neighborhood averaged over genotypes (column 2, green = depleted; purple = enriched) and scaled expression of either aggregated expression of vulnerability or resilience genes (columns 3 and 4), or expression of the top 5 most strongly correlated individual protein-coding genes (columns 5-14, red = high; blue = low). **J-K:** Relationship between average expression of vulnerability-associated (J) or resilience-associated (K) genes and neighborhood-level DA across excitatory neuron populations. **L:** Gene set enrichment analysis (GSEA) along the continuum of vulnerability- and resilience-associated genes identified enrichment of oxidative phosphorylation, respiratory chain, ATP synthesis, synaptic assembly, and neuronal structural programs in vulnerable neurons under normal conditions. The most highly significant GO terms (FDR < 0.1) were hierarchically clustered and collapsed. **M:** Resilience-associated gene programs include positive regulation of mRNA catabolic processes (NES = 1.76; adj. P = 0.003), whereas vulnerability-associated programs included respiratory chain complex pathways (NES = 1.77; adj. P = 0.002).

Non-neuronal cells were largely unaffected at the subtype level, but distinct subsets of astrocytes showed enrichment and depletion in sporadic FTD & FTD-MND. Further analysis revealed a total of four molecularly distinct astrocyte classes demarcated by expression of *RERG*, *SLC38A1*, *CHI3L1* and *SLC24A2* (Extended Data Fig. 8a-b). The most depleted astrocyte class (subcluster 2) highly expressed markers of reactive astrocytes^55^, *CHI3L1*, *NDRG1* and *SERPING1* (Extended Data Fig. 8c-d), while a relatively enriched astrocyte class (subcluster 1) highly expressed *JUNB*, *ID2* and *ID3* (Extended Data Fig. 8e), corresponding to the Ast *Serpinf1* astrocyte class in the mouse neocortex^56^.

Patterns of cellular neighborhood depletion and enrichment were highly correlated across genotypes for the most depleted cell types (Pearson’s r = 0.87) (Fig. 3c), and among ENs overall (Pearson’s r = 0.84) (Fig. 3d). Patterns of neighborhood depletion were also robust across sorting conditions (Extended data. Fig. 9B-E), and limited to FTD clinical groups (FTD and FTD-MND), consistent with the known regional vulnerability profile of MND (Extended data. Fig. 9H-J). Because the same control samples were used to reference each of the disease subtypes, this correlation could be inflated by shared variation in control measurements across comparisons. Therefore, we further evaluated correlations within cell types by permuting control samples such that independent controls were compared to each genotype. Across permutations, strong correlations in differential abundance estimation persisted across all cell types (Extended data. Fig 10a), further supporting the shared cell type vulnerabilities observed in *C9orf72* and sporadic FTD.

We next explored which aspects of neuronal identity relate to FTD-related vulnerability vs. resistance. L5 ET, L5 IT NOX4, L2/3 IT FREM3 and L2/3 IT ANRKD30A were the most differentially abundant populations in disease (Fig. 3b & Suppl. Table 3a) and contained large nuclei (Extended Data Fig. 1h). As large neurons are reported to be vulnerable in other neurodegenerative diseases^57,58^ and nuclear size correlates with cell body size^59^, we further examined whether neighborhoods with higher proportions of large nuclei in control samples from the LN10 sort showed increased vulnerability in patients. Across all ENs, we observed that greater neighborhood-level size enrichment predicted lower abundance in disease (Pearson’s r = -0.29; p<0.05 in *C9orf72*, and -0.20; p<0.05 in sporadic) (Fig. 3e). A particularly strong negative correlation between the neighborhood nucleus size enrichment and abundance in disease was observed within the L2/3 IT ANKDR30A population (Pearson’s r = -0.83; p<0.05 in *C9orf72*, and -0.87; p<0.05 in sporadic) (Fig. 3e-f). For L5 IT NOX4, L5 ET and L2/3 IT FREM3, the other strongly depleted cell types, this correlation was weaker (Fig. 3f; Extended Data Fig. 10b), suggesting that factors other than size confer vulnerability.

Our prior work has shown that nuclear and soma size are increased in patient VENs and fork cells lacking TDP-43 pathobiology^8^. In contrast, when these neurons exhibit nuclear TDP-43 depletion, with or without an accompanying cytoplasmic inclusion, they undergo striking nuclear and somatodendritic atrophy. Similarly, the size of disease-vulnerable spinal motor neurons was shown to increase prior to subsequent degeneration in an ALS mouse model^60^ and following axonal damage^61^. While these findings suggest that neuronal enlargement represents an early step in the pathogenic cascade preceding TDP-43 pathobiology, the extent to which this property generalizes across cell types remains unknown. Thus, we sought to examine whether neuronal size differences in disease were also captured in our data by comparing the LN10 sort enrichment scores in control samples with those in patients (Methods). Using LN10 enrichment as a proxy for size, we found that excitatory neurons are enlarged in disease, while inhibitory neurons are not (Fig. 3g). We expanded this analysis to the broader cell taxonomy and found broad LN10 enrichment across EN subtypes and cortical layers (Extended Data Fig. 11a-b). Interestingly, this approach did not reveal size enrichment in disease for the largest (L2/3 IT ANKDR30A, L5 ET and L5 IT NOX4) and smallest (L4 IT ALDH1A1) cell classes (Extended Data Fig. 11c-d), likely because these classes are almost exclusively found in one sorting condition (Fig.1j), limiting the dynamic range of LN10 logFC values. Alternatively, nuclear shrinkage as a consequence of TDP-43 pathobiology could also explain this finding, especially considering that nuclear enlargement proved most difficult to detect in the most vulnerable neuron types.

To identify molecular features associated with vulnerability, we next leveraged these neighborhood-level depletion patterns to relate selective neuronal loss to transcriptional states in control neurons. Specifically, we mapped differential abundance values observed in disease onto corresponding neighborhoods and quantified the relationship between baseline gene expression and FTD-related depletion in control cells within these neighborhoods. This analysis revealed genes whose expression in control neurons was strongly and convergently associated with vulnerability across neighborhoods (Fig. 3h) (Methods). Higher expression of several genes predicted vulnerability in bvFTD, including *RGS4*, *NRGN*, *MAST4* and *FGF9*, while additional genes including the anti-apoptosis gene, *CFLAR*^62^, and the DNA damage repair gene, *RAD51*^63^, predicted resilience (Fig. 3h). Aggregating these signals across genes identified coherent transcriptional programs associated with vulnerability and resilience (Fig. 3i-k), with gene set enrichment analysis highlighting cellular respiration, ATP synthesis, and proton transport pathways among the strongest correlates of vulnerability (Fig. 3l). Strikingly, neighborhoods even within each given neuronal subtype displayed correlations of expression of vulnerability- or resilience-associated genes with depletion (Fig. 3j-k). Conversely, programs related to mRNA catabolism were associated with relative resilience (Fig. 3m). Together, these findings identify molecularly defined cell types that are vulnerable in disease and uncover genetic programs that are shared across these vulnerable cell types, pointing towards underlying features of selective neuronal vulnerability and resilience in FTD.

### Differential vulnerability across L5 ET neuron subtypes

VENs and fork cells represent the best documented selectively vulnerable neuron types in bvFTD^10,16,64^. Their extratelencephalic projection neuron identity was suggested by expression of *FEZF2 and CTIP2*^65^ and confirmed within a modern taxonomic framework ^66^. Several additional VEN/fork cell markers have been identified, including *VMAT2, GABRQ,* and *ADRA1A*^67^; *VAT1L*, *CHST8*, *LYPD1*, and *SULF2*^68^; *POU3F1, BMP3, ITGA4*^66^, and collagen forming genes *COL5A2*, *COL24A1*, and *COL21A1*^69^. Recent studies suggest, however, that these markers may reflect broad lineage identity rather than distinctly marking VENs and fork cells^70^. Indeed, most of these markers are expressed across L5 ET neurons irrespective of morphotype (Extended Data Fig. 12a). Although clear molecular distinctions among L5 ET subtypes have been difficult to identify due to the scarcity of this subpopulation, the 7.5-fold enrichment of L5 ET neurons we achieved by sorting for large nuclei (Fig. 1c) enabled us to explore L5 ET molecular heterogeneity and how it relates to neuronal loss in bvFTD.

Iterative clustering of L5 ET neurons revealed three major clusters: L5 ET CSN1S1, L5 ET TMPRSS15, and L5 ET TOX3 (Fig. 4b). Gene regulatory network analysis via SCENIC^71^ further highlighted upstream factors that may control differences among these populations. Specifically, L5 ET CSN1S1 was distinguished by gene regulatory networks linked to POU3F1, HMGB1, BHLHE40, MEF2D, KLF9, JUN, metabolic regulators PPARGC1A and ESRRG, early response genes JUND and EGR3, and oxidative stress response gene NFE2L1 (Fig. 4c & Extended Data Fig. 13a). L5 ET TMPRSS15 was distinguished by PBX3, STAT4, POU2F1, and TCF4, consistent with transcriptional programs involved in neuronal identity maintenance and responsiveness to extracellular signaling pathways (Fig. 4c & Extended Data Fig. 13b). L5 ET TOX3 was distinguished by CNOT4 and TAF1, factors involved in mRNA turnover and transcriptional initiation, respectively, and shared many marker genes with L5 ET CSN1S1 that were less highly expressed in the L5 ET TMPRSS15 class, such as *CREM*, *NFAT5*, *CREB5* and *BACH2*, indicating high homology between these two classes and the relatively distinct transcriptional nature of the TMPRSS15 class (Fig. 4c & Extended Data Fig. 13c).

**Fig. 4.**
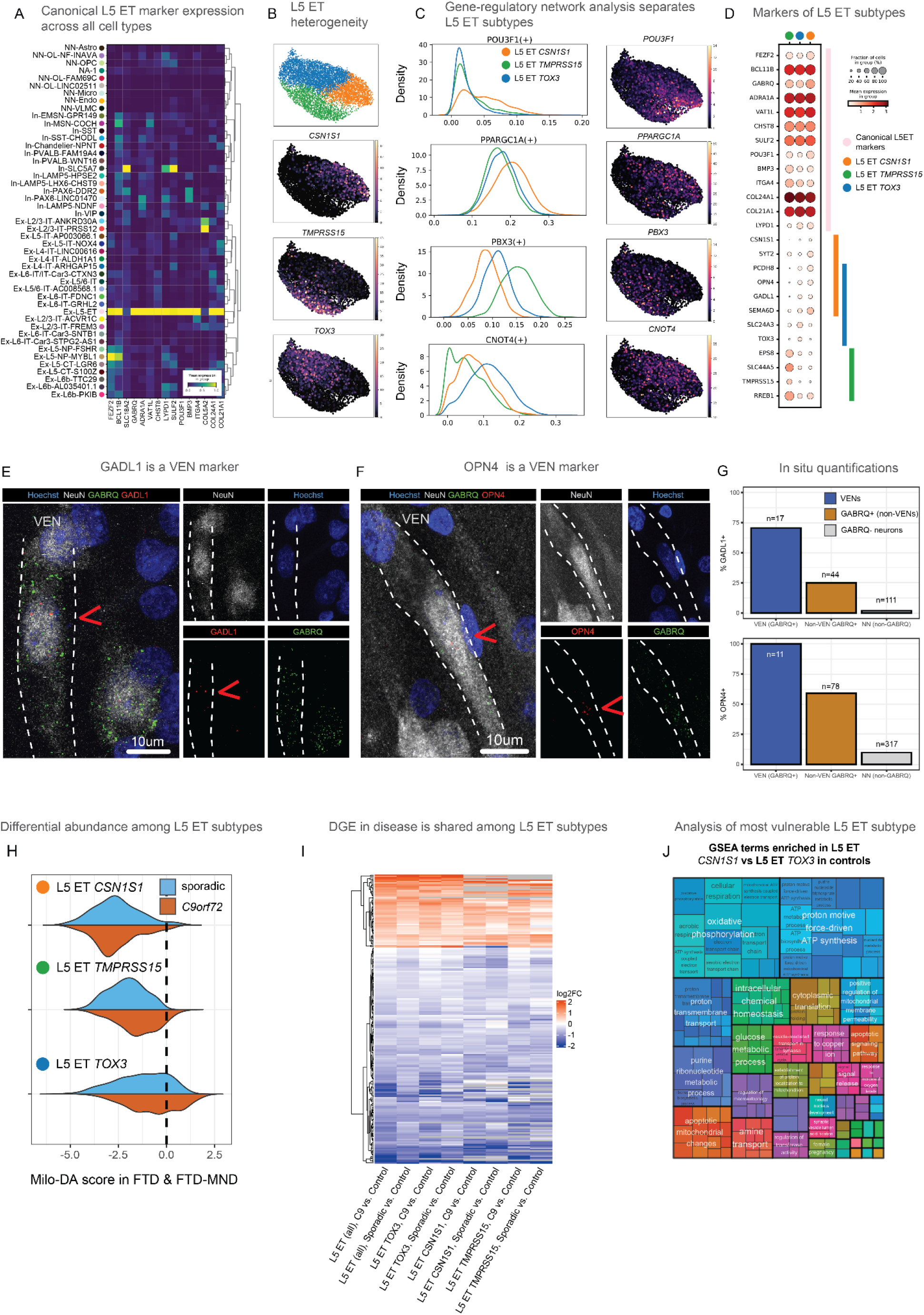
Molecularly distinct L5 ET subclasses exhibit differential vulnerability in bvFTD. **A:** Expression of canonical L5 ET/VEN-associated marker genes across transcriptionally defined FI cell populations demonstrates a distinct L5 ET population within the FI dataset. **B:** Subclustering analysis of L5 ET neurons identified three molecularly distinct subclasses characterized by expression of *CSN1S1* (L5 ET *CSN1S1*), *TOX3* (L5 ET *TOX3*), and *TMPRSS15* (L5 ET *TMPRSS15*). **C:** Gene regulatory network analysis (SCENIC) identified distinct regulon enrichment patterns across L5 ET subclasses. L5 ET CSN1S1 neurons were distinguished by gene regulatory networks linked to *POU3F1*, and metabolic regulator *PPARGC1A*. L5 ET TMPRSS15 was distinguished by *PBX3*. L5 ET TOX3 was distinguished by *CNOT4,* a factor involved in post-transcriptional regulation. Additional regulon hits are shown in Ext. Data Fig. 13. **D:** Expression of canonical L5 ET/VEN-associated markers and subtype-associated markers across L5 ET subclasses. L5 ET *TMPRSS15* neurons highly expressed *TMPRSS15*, *SLC44A5*, and *EPS8*, whereas L5 ET *TOX3* neurons were enriched for *TOX3* and *SLC24A3*. L5 ET *CSN1S1* neurons highly expressed *CSN1S1* and *SYT2*. *GADL1*, *OPN4*, and *PCDH8* expression was shared between L5 ET *TOX3* and L5 ET *CSN1S1* neurons. **E:** *GADL1,* highly expressed in L5 ET *CSN1S1* and L5 ET *TOX3* based on transcriptional analysis (Extended Data Fig. 12b) was validated in situ using RNAscope (Methods), revealing *GADL1* expression in 70.1% (12/17) of morphologically defined VENs, 25% of non-VEN *GABRQ*+ neurons (11/44) and 1.8% of *GABRQ*- neurons (2/111), thus accurately differentiating VENs from other GABRQ expressing L5 ET cells. **F:** *OPN4,* also highly expressed in L5 ET *CSN1S1* and L5 ET *TOX3* based on transcriptional analysis (Extended Data Fig. 12b), was validated in situ using RNAscope revealing expression in 100% of morphologically defined VENs (11/11), 59% of non-VEN *GABRQ*+ neurons (46/78) and 9.8% of *GABRQ*- neurons (31/317), thus differentiating VENs from other *GABRQ*-expressing L5 ET cells. **G:** Quantifications of the proportion of morphologically defined VENs, non-VEN *GABRQ*+ neurons and *GABRQ*- neurons that express *GADL1* and *OPN4* as described in panels E and F. **H:** Neighborhood-level *Milo* differential abundance analysis among the three molecularly defined L5 ET classes revealed the strongest depletion among L5 ET *CSN1S1* cells with the lowest depletion among L5 ET *TOX3* cells. Based on expressions of *GADL1* and *OPN4* which were validated to be VEN markers in situ, these two classes contain the most VENs among the three L5 ET subclasses. **I:** Differential gene expression analysis across L5 ET subclasses in *C9orf72* and sporadic FTD & FTD-MND. Disease-associated transcriptional responses were broadly shared across L5 ET subclasses and genotypes. **J:** Gene set enrichment analysis of DEGs between control L5 ET *CSN1S1* and L5 ET *TOX3* neurons. The more vulnerable L5 ET CSN1S1 subtype was enriched for oxidative phosphorylation, respiratory chain, ATP synthesis, and mitochondrial pathways.

L5 ET CSN1S1 was further transcriptionally characterized by expression of *CSN1S1, GADL1, OPN4, PCDH8* and *SYT2* (Fig. 4d & Extended Data Fig. 12b). L5 ET TOX3 shared moderate expression of *GADL1* and *OPN4*, with enrichment for *SLC24A3* and *TOX3*. In contrast, L5 ET TMPRSS15 lacked expression for these markers, but showed specific expression of *TMPRSS15* and enrichment for *SLC44A5, EPS8 and RREB1* (Fig. 4b & Extended Data Fig. 12b). The expression of previously nominated VEN markers also varied across L5 ET subtypes, with *LYPD1*, identified via RNAseq of laser-microdissected VENs^68^, enriched in L5 ET CSN1S1 and L5 ET TOX3 clusters, while *FEZF2*, *GABRQ* and *ADRA1A* showed broad expression across all three subclusters (Fig. 4d & Extended Data Fig. 12a).

To link molecularly defined L5 ET subtypes to specific L5 ET morphotypes, we performed RNAscope fluorescent in situ hybridization for L5 ET subtype candidate genes, selecting from the genes most differentially expressed in the three clusters. We co-labeled with NeuN (protein immunofluorescence) and *GABRQ* (RNA) to broadly label the L5 ET class. *GADL1* was expressed in 70.1% (12/17) of morphologically defined VENs, 25% of non-VEN *GABRQ*+ neurons (11/44) and 1.8% of *GABRQ*- neurons (2/111)(Fig. 4e,g). *OPN4*, on the other hand, was expressed in 100% of morphologically-defined VENs (11/11), but also a higher proportion of non-VEN *GABRQ*+ neurons (59%, 46/78) and *GABRQ*- neurons (9.8%, 31/317) (Fig. 4f,g).

By contrast, other tested L5 ET subtype markers did not appear to discriminate between VEN and non-VEN L5 ETs. *RREB1* was expressed in 93% of VENs (13/14), 84% of non-VEN *GABRQ*+ neurons (90/107), and 15% of *GABRQ*- neurons (49/321) (Extended Data Fig. 14a-c). Additionally, *TMPRSS15* was expressed in 38% of VENs (14/37), 41% of non-VEN *GABRQ*+ neurons (71/174), and 23% of *GABRQ*- neurons (136/585) (Extended Data Fig. 14d-f). These RNAscope results are consistent with the transcriptional continuum observed among L5 ET populations in our data (Extended Data Fig. 12a-b) and suggest the VEN morphology is distributed across closely related transcriptional states, with GADL1 and OPN4 being more prevalent in VENs yet not rising to the level of a morphotype-specific marker. Combined, these observations suggest that most morphologically defined VENs are found in the L5 ET CSN1S1 and L5 ET TOX3 transcriptional clusters.

Having refined the L5 ET neuron subtypes, we next asked whether they differed in their vulnerability in FTD. The L5 ET CSN1S1 cluster showed the strongest depletion in both *C9orf72* and sporadic FTD (Fig. 4h), suggesting that the reduction of *CSN1S1* expression observed in FTD across ENs (Fig. 2c) may have been primarily driven by this L5 ET subcluster since *CSN1S1* is lowly expressed in other EN clusters. To examine whether differential L5 ET subtype vulnerability relates to transcriptional consequences of disease, we used *Dreamlet* to identify DEGs in patients compared to controls in each L5 ET subtype and genotype. Among the cells still present in patients, the direction of effect and magnitude of DEGs were largely shared between L5 ET subtypes and genotypes (Fig. 4i), consistent with findings across EN populations (Fig. 2b-c), with minor differences in significant DEGs and gene programs between L5 ET subtypes (Extended Data Fig. 15a-d).

To further examine molecular properties linked to L5 ET subtype vulnerability, we compared gene expression between subtypes in healthy controls. We focused on differences between L5 ET CSN1S1, which showed the greatest depletion among L5 ET subtypes, and L5 ET TOX3 neurons, which showed the least depletion. GSEA on the resulting DEGs highlighted enrichment of oxidative phosphorylation pathways in the more vulnerable L5 ET CSN1S1 class (Fig. 4j), and complementary over-representation analysis on the same genes revealed significant enrichment of oxidative phosphorylation and electron transport gene programs in L5 ET CSN1S1 cells, with enrichment of centriole gene programs in the L5 ET TOX3 class (Extended Data Fig. 15e).

Collectively, these results reveal previously unappreciated molecular heterogeneity among L5 ETs that corresponds to morphological distinctions and differential vulnerability linked to higher baseline oxidative phosphorylation and bioenergetic demand.

## Discussion

Selective neuronal vulnerability is a defining feature of neurodegenerative disease^72^, yet the properties that distinguish vulnerable from resilient neurons in each disease remain unclear. Here, we identified a striking contrast between widespread, convergent transcriptional responses to disease and highly selective patterns of neuronal loss across molecularly defined cell types. While excitatory neurons broadly upregulate dendritic, axonal, and synaptic gene programs and downregulate cilia-related pathways in *C9orf72*-associated and sporadic disease, only specific subpopulations—particularly within L2/3 and L5 projection neuron classes—undergo depletion. By linking these patterns of selective neuronal loss to transcriptional signatures observed in healthy control neurons, we identified baseline molecular programs associated with vulnerability. These findings support a framework in which vulnerability emerges, at least in part, from cellular identity and state rather than a cell type-specific transcriptional response to disease.

Vulnerable cell classes comprise highly specialized populations, including L5 ET and subsets of L2/3 and L5 IT neurons with features of large, long-range projection neurons. Within these populations, vulnerability is further stratified by baseline metabolic state, with elevated expression of cellular respiration and ATP synthesis pathways associated with greater vulnerability. Notably, shared transcriptional responses in disease do not distinguish vulnerable from resilient classes, suggesting they may reflect compensatory remodeling, network stress, or maladaptive plasticity rather than upstream drivers of selective degeneration. These findings are consistent with a model in which neurons with high structural and bioenergetic demands are preferentially susceptible to degeneration in FTLD/MND-TDP. These metabolic state differences may intersect with other cellular features early in the disease process, including increased cell size^8^ and hyperexcitability, the latter shown to promote alternative splicing and depletion of nuclear TDP-43 in cultured neurons^73,74^, linking physiological stress to TDP-43 pathobiology. Remarkably, a recent electrophysiological study of *ex vivo* human VENs showed evidence for heightened excitability and longer action potential duration compared to other L5 ET neurons^70^. Collectively, these observations suggest that heightened metabolic and functional burden may i) promote TDP-43 pathobiology or ii) increase vulnerability to its downstream consequences, thereby focalizing early disease within the largest, most physiologically active projection neurons.

Histopathological studies spanning decades have consistently shown L5 ETs, and particularly their specialized morphotypes (VENs, fork cells, and Betz cells), to develop the earliest TDP-43 pathology and undergo selective degeneration in bvFTD and ALS^3–9^. Single-cell studies, while previously limited in their recovery of these rare cell types, have provided unbiased support for the claim that L5 ETs are particularly vulnerable, when vulnerability is defined by transcriptional dysregulation^6^, cryptic exon burden^15^, and robust depletion in disease^6^. Leveraging a uniquely large L5 ET sample enabled by size-based FANS, we identified molecular heterogeneity that newly refines the L5 ET class. By resolving distinct transcriptional subtypes, we show that L5 ET vulnerability is not uniform, as previously suggested^53^, but is accentuated in specific subpopulations characterized by elevated expression of oxidative phosphorylation and electron transport pathways and enriched for genes expressed by VENs in FI. These observations further link baseline identity and metabolic state to selective vulnerability at a finer scale and provide a candidate bioenergetic mechanism for the selective depletion of morphologically specialized L5 ETs in bvFTD and ALS.

Recent studies have also explored the molecular landscape of the FTD-ALS spectrum using snRNA-seq. Consistent with our findings, multiple studies demonstrated convergent transcriptional changes in disease, spanning genotype, neuron type, brain region, and disease subtype^6,14^. Critically, however, compared to previous literature, our study refines our understanding of neuron type-specific loss in bvFTD and its transcriptional underpinnings. Prior studies either did not assess compositional changes^6^, could not detect them^13,14^, or identified cell type depletions limited to a single clinical subtype, genotype, or brain region^15,75^. Likely due to limited and variable sampling of vulnerable cell types, these largely null findings contrast with histological observations and have led to alternative definitions of vulnerability based on the magnitude of transcriptional alteration in disease^6^. By contrast, our FANS-based enrichment for vulnerable neuron types and neighborhood-level compositional approach uncovered several molecularly defined vulnerable subtypes (L5 ET, L2/3 IT, and L5 IT), convergent across *C9orf72* and sporadic bvFTD. These findings reconcile single-cell data with longstanding histological observations and, taken together with previous studies, reinforce the notion of a broadly shared transcriptional response to disease that can be decoupled from the underlying factors that render specific neuron types most vulnerable to degeneration.

Several limitations should be considered. First, postmortem human data is inherently cross-sectional and correlative in nature. Thus, our study identifies associations between molecular cell identity and vulnerability rather than causal links, and it remains unclear whether disease-associated gene expression changes reflect dysfunction upstream or downstream of TDP-43 pathology or broader neurodegenerative and neuroinflammatory responses. Second, our enrichment strategy leverages sorting for the largest ∼10% of neuronal nuclei as a proxy for neuronal size, providing high resolution among large projection neuron populations but limiting sensitivity for neuronal classes composed of smaller neurons, including many inhibitory neuron types. As a result, size-dependent relationships with vulnerability may be underestimated or unresolved in these populations. We mitigated this issue by including the unbiased Hoechst+ condition, and previous snRNAseq and histological studies have failed to implicate smaller interneuron populations despite the lack of size bias. Finally, our focus on the frontoinsular cortex provides high-resolution insight into vulnerability in this key, early-affected region but may not capture the full spectrum of cell type-specific degeneration across other cortical and subcortical or limbic areas.

## Supporting information

Suppl.Table_1_Documentation

Suppl.Table_3_Documentation

Suppl.Table_4_Documentation

Suppl_Table_2_Documentation

## Data availability

Raw sequencing data will be available on dbGaP as of the date of publication. Note that several control donors did not consent to data sharing and raw data will not be able to be made public. The full integrated dataset has been compiled into an interactive UCSC Cell Browser (https://cells-test.gi.ucsc.edu/?ds=frontoinsular-ctx-als-ftd). Code necessary to reproduce the analyses is available on GitHub: https://github.com/Pollen-lab/Transcriptional_Atlas_Frontoinsular_Cortex_in_ALS_and_FTD.git. Any additional information required to reanalyze the data reported in this paper is available from the lead contact upon request.

## Acknowledgments

We thank Donor Network West and all tissue donors and their families for their generous contributions to research. We thank Dan Mordes and members of the Pollen and Seeley laboratories for comments and intellectual discussions, Dmitry Velmeshev for sharing his nuclei isolation protocol, Matthew Schmitz for assistance with cell type interpretation, David Aley for help setting up the Milo analysis, Nathan Schaefer for developing CellBouncer, Jane Srivastava and the Gladstone Flow Cytometry Core, and Sarah Elmes and the Helen Diller Cancer Center Flow Cytometry Core. We also thank the patients and their families for their invaluable contributions to FTD/MND research. Sequencing was performed at the UCSF CAT, supported by UCSF PBBR, RRP IMIA, and NIH grant S10OD028511-01. This study was supported by R01NS104437 and a Genentech research award to W.W.S. and A.A.P as well as by R01AG087959, the Pershing Square Foundation, and Schmidt Futures Foundation to A.A.P.; A.A.P. is a New York Stem Cell Foundation Robertson Investigator. Additional support was provided by Schmidt Science Fellows and Jane Coffin Childs fellowships to JLW, NIH grants AG019724, AG062422, AG063911, and AG057195, the Rainwater Charitable Foundation, and the Bluefield Project to Cure FTD to W.W.S.

## Author contributions

A.B., A.A.P., and W.W.S. conceived the project and experimental design. A.A.P., and W.W.S. supervised the study and secured the funding. A.N. curated all tissue samples under supervision of W.W.S. Experiments were performed by A.B. Data was analyzed by A.B., D.I., and L.H-P. with help from J.L.W. and J.W.D under the supervision of A.A.P. and W.W.S. Initial brainstorming and piloting of reference mapping was done with help from F.L.P. RNAscope was performed by S.V. Confocal microscopy was done by A.B. Confocal images were analyzed and quantified in a blinded manner by A.B. and A.T.M. Histological verification of morphological VEN identity was done by S.V. Patient metadata was curated by K.F. Pathological analysis of all cases were performed by S.S., L.T.G., and W.W.S. J.S.Y provided genotyping support, and M.G.T., H.J.R., and B.L.M. provided funding support and clinical evaluations. F.M.J.J provided A.B. support with implementing the manuscript into a PhD dissertation. The manuscript was prepared by A.A.P., W.W.S., A.B., L.H-P., and D.I. with input from all authors.

---

**Extended Data Fig. 1.**
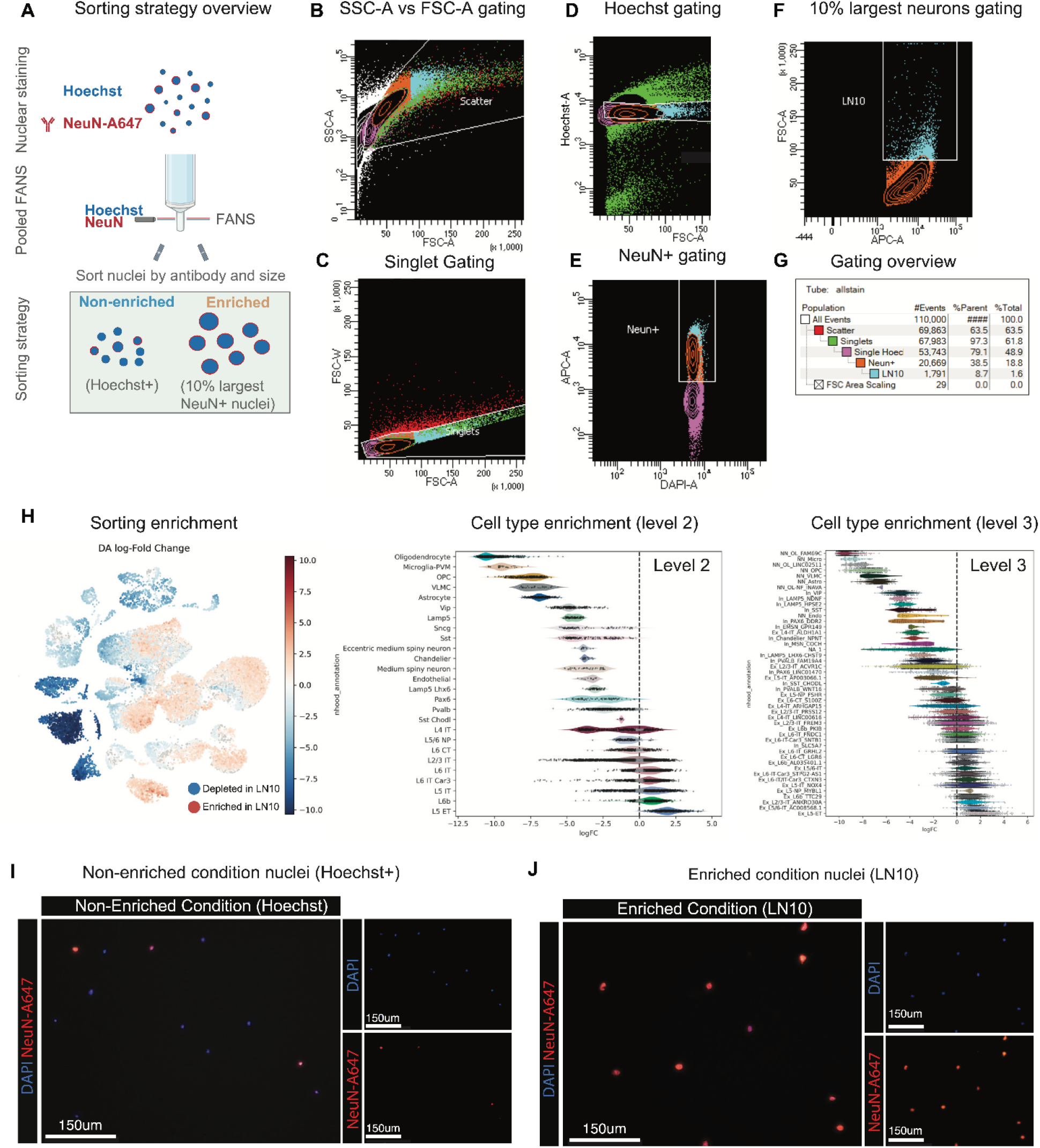
Size-based FANS enrichment of large neuronal nuclei increases recovery of vulnerable neuronal subclasses. **A:** To enrich the highly vulnerable L5 ET neuron class, nuclei were stained with Hoechst and NeuN-A647. Nuclei were next gated into a non-enriched Hoechst+ group and a size-gated enriched group consisting of the largest 10% of neurons (LN10). **B:** Debris was excluded by gating nuclei based on SSC-A and FSC-A signals. **C:** Putative single nuclei were gated using FSC-W and FSC-A signals. **D:** Single nuclei were gated by gating a single Hoechst+ population. **E:** Neuronal nuclei were identified by NeuN-A647 signal. **F:** Large neuronal nuclei were enriched by gating the ∼10% largest neuronal nuclei (LN10) based on FSC-A signal. **G:** Our gating strategy captured both a non-enriched Hoechst+ condition and an enriched LN10 condition consisting of ∼10% of the largest neurons. **H:** Differential abundance analysis comparing non-enriched and LN10-enriched conditions demonstrated preferential enrichment of large neuronal subclasses in the LN10 sort. Milo differential abundance results are shown for both level 2 and level 3 cell taxonomies. Several transcriptionally distinct neuronal subclasses displayed differential enrichment, including enrichment of L5 ET-NOX4 and depletion of related sister populations such as L5 IT-AP003066.1. Size-based enrichment additionally resolved subtype-specific distinctions within L4 IT, L5 NP, and L5 CT populations. **I-J:** Post-sorting validation of nuclei from the two sorting conditions showed that the non-enriched condition consisted of both neurons and non-neurons including nuclei of neurons with a smaller nuclear diameter, while the enriched condition exclusively consisted of neurons with a large nuclear diameter.

**Extended Data Fig. 2.**
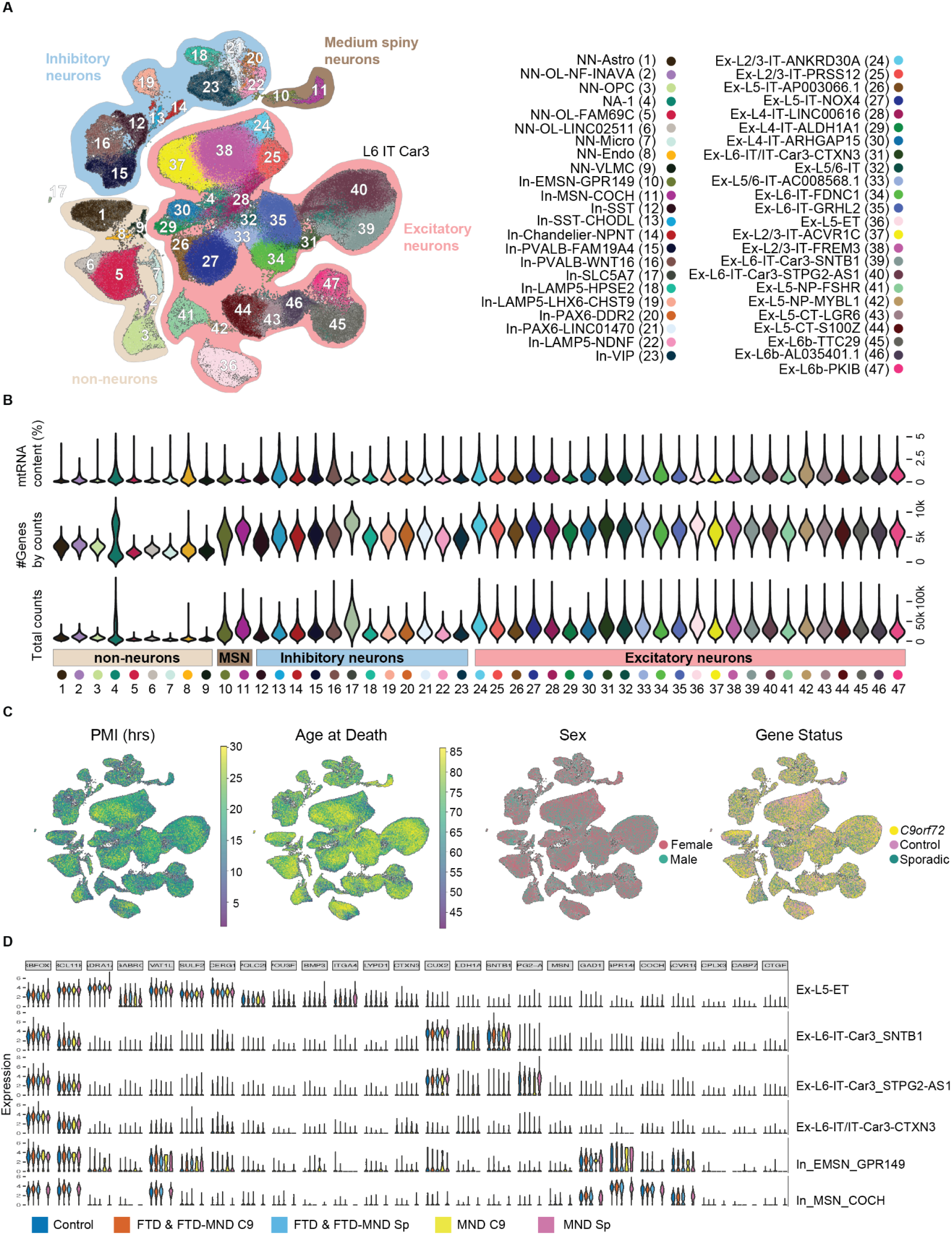
Integrated snRNA-seq analysis resolves diverse FI neuronal subclasses and enriches excitatory neuron diversity. **A:** UMAP visualization of the integrated snRNA-seq dataset showing the most granular taxonomy (cell level 3) comprising 47 transcriptionally defined cell populations, including excitatory neurons, inhibitory neurons, medium spiny neurons, and non-neuronal cell types. Large-neuron enrichment (LN10) increased resolution of excitatory neuron subclasses. **B:** Distribution of quality-control metrics across the 47 molecularly defined cell populations, including mitochondrial RNA (mtRNA) content, number of detected genes, and total transcript counts. **C:** UMAP projections colored by postmortem interval (PMI), age at death, sex, and gene status, demonstrating robust integration across technical and biological covariates. **D:** Expression of representative marker genes across major neuronal subclasses and disease groups validates transcriptional identities of molecularly defined cell populations. A subset of representative populations is shown.

**Extended Data Fig. 3.**
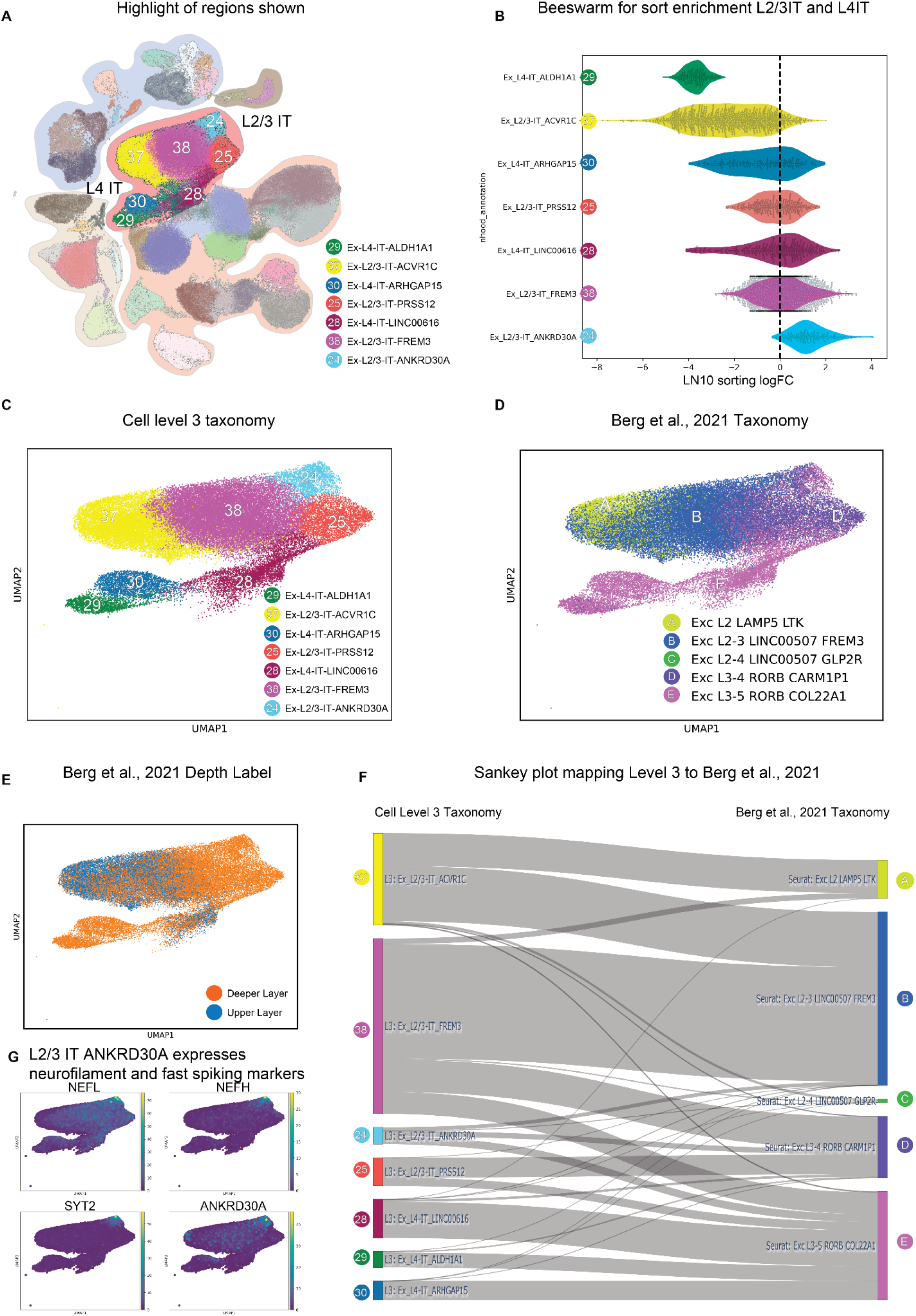
Frontoinsular cortex L2/3 IT populations exhibit size- and depth-associated correspondence with middle temporal gyrus excitatory neuron classes. **A:** UMAP highlighting the L2/3 IT and L4 IT excitatory neuron populations examined in subsequent analyses. **B:** Differential abundance analysis of LN10 enrichment across L2/3 IT and L4 IT subclasses. Positive logFC values indicate enrichment in the LN10 sort, whereas negative values indicate depletion. L2/3 IT ANKRD30A cells were strongly enriched, whereas L2/3 IT ACVR1C cells were strongly depleted. **C:** Cell level 3 taxonomy of FI L2/3 IT and L4 IT excitatory neuron populations identified in this dataset. **D:** Reference mapping of FI excitatory neuron populations onto the Berg et al., 2021 middle temporal gyrus (MTG) excitatory neuron taxonomy. **E:** Upper- versus deeper-layer classification of Berg et al., 2021 reference populations, showing upper and deeper-layer excitatory neuron identities for the different L2/3 IT and L4 IT excitatory neuron populations. For example, L2/3 IT ACVR1C maps onto upper layer neurons from Berg et al., 2021, while L2/3 IT ANKDR30A maps onto deeper layer neurons. **F:** Sankey plot showing correspondence between FI level 3 excitatory neuron subclasses and Berg et al., 2021 MTG excitatory neuron populations. **G:** The L2/3 IT ANKRD30A class highly expresses neurofilament genes NEFL and NEFH, as well as the synaptic protein SYT2.

**Extended Data Fig. 4.**
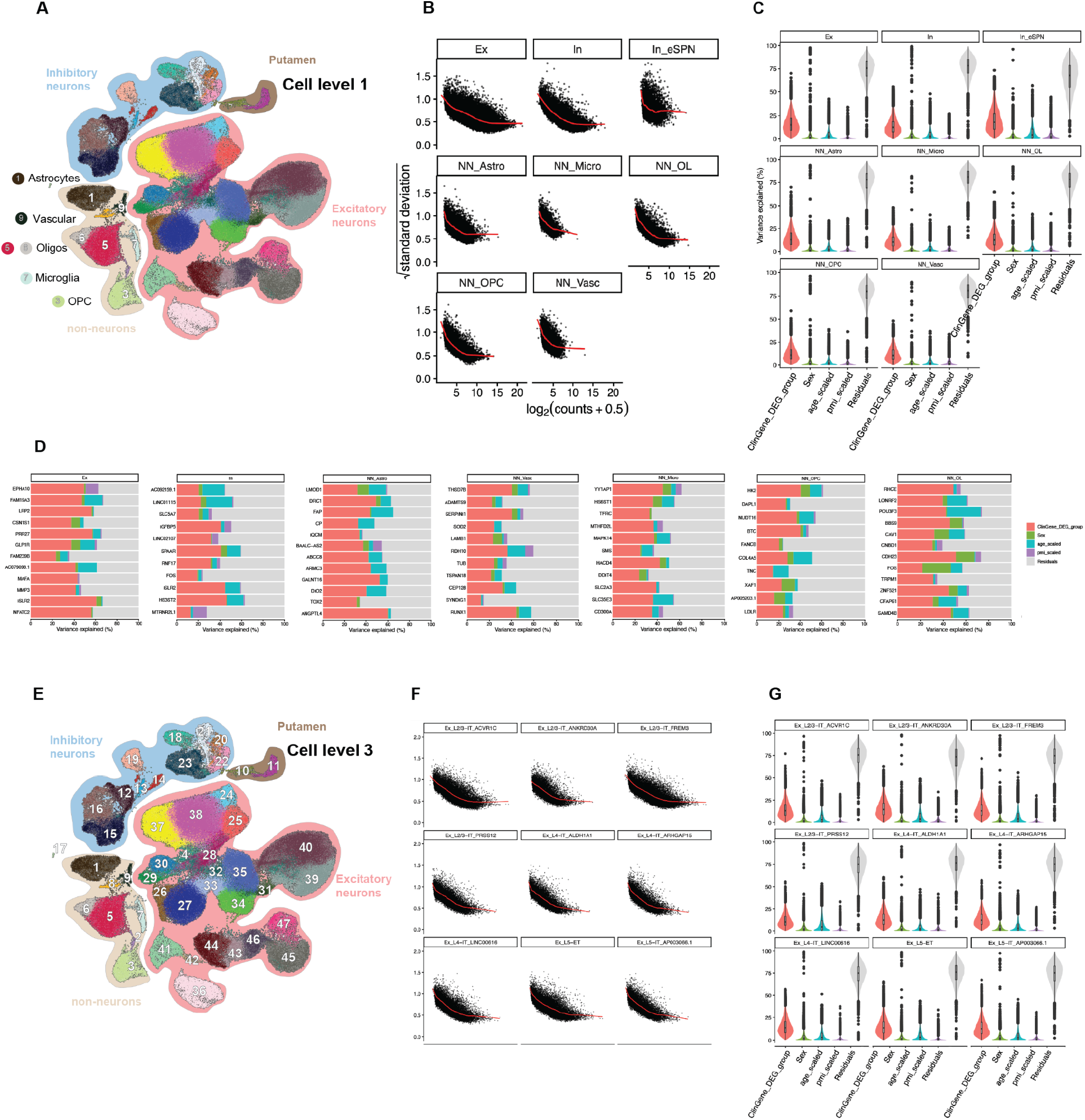
Dreamlet modeling and variance partitioning across FI cell populations. **A:** UMAP showing the broad cell class taxonomy (cell level 1) used for Dreamlet pseudobulk differential expression modeling, including excitatory neurons, inhibitory neurons, putamen-derived neurons, astrocytes, vascular cells, oligodendrocytes, microglia, and OPCs. **B:** Mean–variance trend plots (voom) for each broad cell class included in the Dreamlet level 1 model. **C:** Variance partition analysis for the Dreamlet level 1 model showing the proportion of transcriptional variance attributable to clinical diagnosis/gene status (ClinGene_DEG_group), sex, age, postmortem interval (PMI), pseudobulk identity, and residual effects across broad cell classes. **D:** Gene-level variance partition analysis for representative broad cell classes. Bars indicate the proportion of variance attributable to clinical diagnosis/gene status, sex, age, PMI, pseudobulk identity, and residual effects for highly variable genes, revealing that a large portion of variance for the most differentially expressed genes is explained by the ClinGene_DEG_group variant. **E:** UMAP showing the fine-grained molecular taxonomy (cell level 3) comprising all 47 transcriptionally defined cell populations used for Dreamlet level 3 differential expression modeling. **F:** Mean–variance trend plots (voom) for representative level 3 cell populations included in the Dreamlet level 3 model. **G**: Variance partition analysis for representative level 3 cell populations showing the proportion of transcriptional variance attributable to clinical diagnosis/gene status (ClinGene_DEG_group), sex, age, PMI, pseudobulk identity, and residual effects, again revealing that a large portion of variance for the most differentially expressed genes is explained by the ClinGene_DEG-group variant.

**Extended Data Fig. 5.**
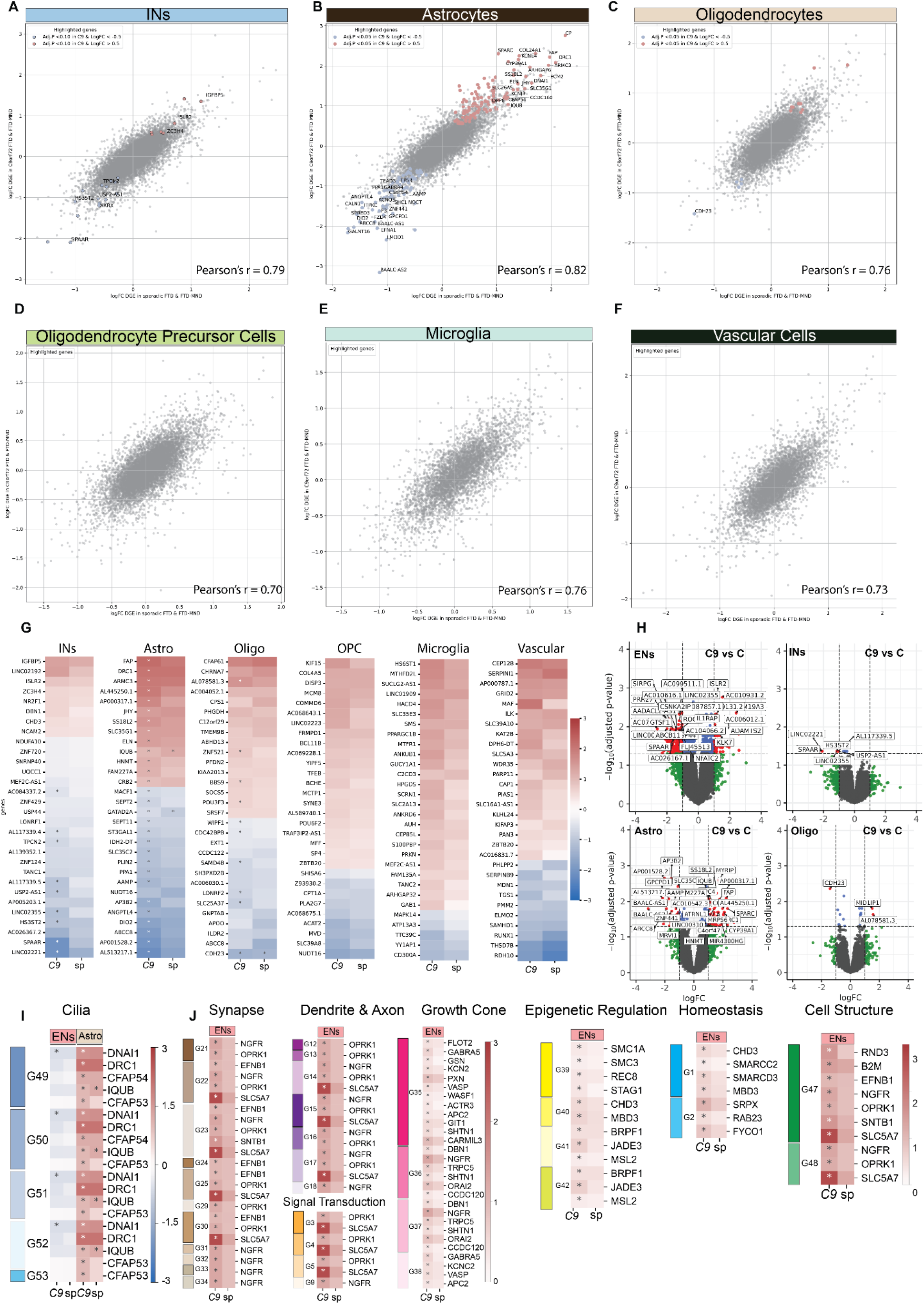
Convergent transcriptional alterations across neuronal and glial populations in *C9orf72* and sporadic FTD & FTD-MND. **A-F:** Comparison of disease-associated differential gene expression (logFC) between *C9orf72* FTD/FTD-MND and sporadic FTD/FTD-MND across inhibitory neurons (A), astrocytes (B), oligodendrocytes (C), oligodendrocyte precursor cells (D), microglia (E), and vascular cells (F). Each point represents a gene. Highlighted genes met significance thresholds in at least one disease group (adj. P < 0.10 for inhibitory neurons and adj. P < 0.05 for all other cell types). Pearson correlation coefficients are shown for each comparison (P < 0.05 for all comparisons shown). **G:** Heatmaps showing representative shared disease-associated transcriptional changes across inhibitory neurons, astrocytes, oligodendrocytes, OPCs, microglia, and vascular cells in *C9orf72* and sporadic FTD/FTD-MND. Shown are genes with the lowest average adjusted P values across genotypes. **H:** Volcano plot highlighting DEGs that pass significance (Adj. P<0.05 & logFC >1 or <1) (red) in *C9orf72* FTD & FTD-MND in ENs, INs, Astrocytes and Oligos. **I:** Representative driver genes contributing to significantly altered pathway programs in FTD, including pathways related to ciliogenesis, synaptic signaling, dendrite and axon structure, growth cone biology, epigenetic regulation, cellular homeostasis, and cell structure (see Fig. 2f; Suppl. Table 2c). For each GO term shown in Fig. 2f, DEGs that pass significance in ENs (adj. P<0.05) in the level 1 Dreamlet analysis (Suppl. Table 1a) are shown.

**Extended Data Fig. 6.**
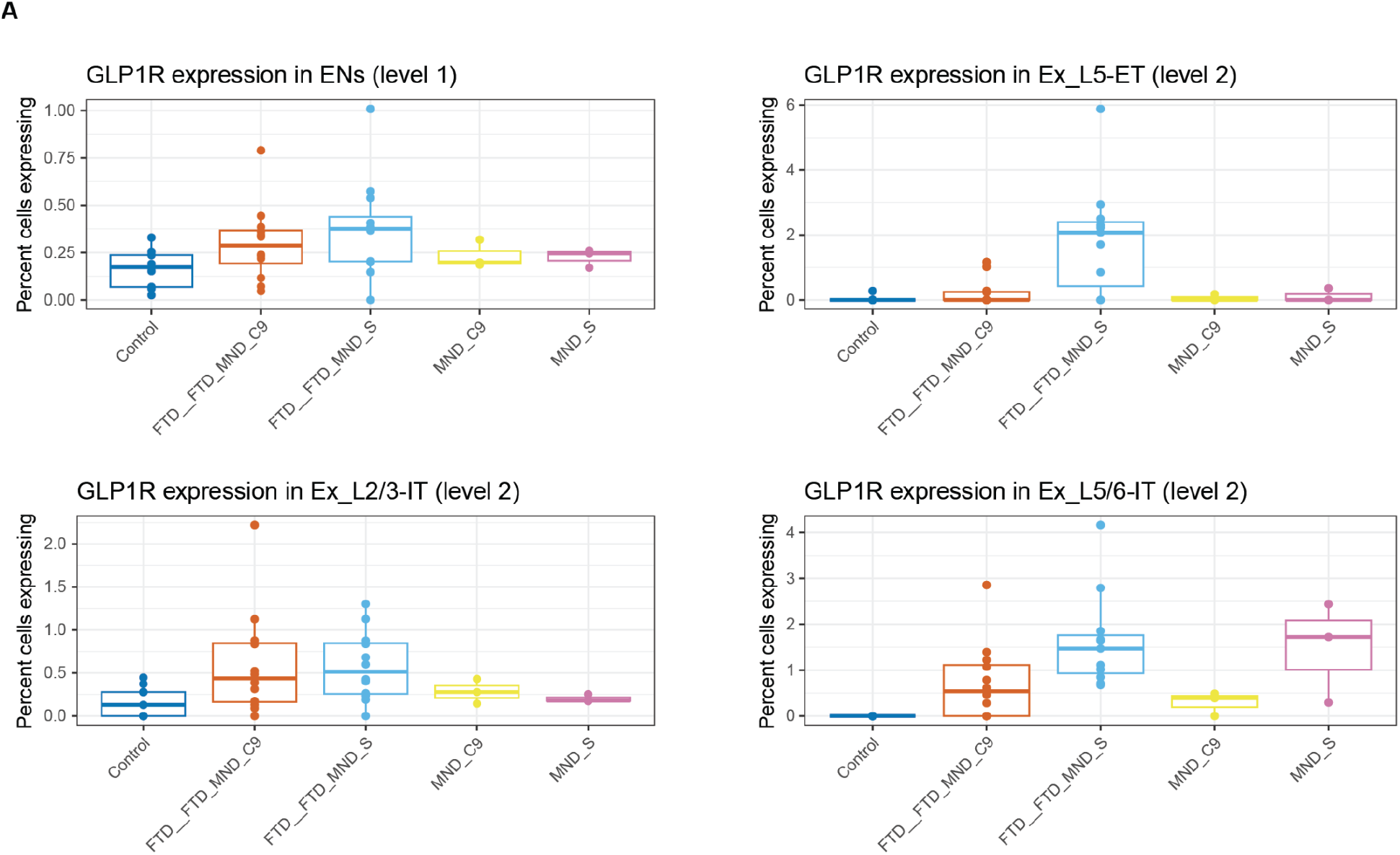
GLP1R-expressing excitatory neurons are increased in FTD/MND. **A:** Percentage of cells expressing GLP1R across excitatory neurons (level 1 taxonomy) and selected excitatory neuron subclasses (level 2 taxonomy), stratified by disease group. Increased GLP1R expression in FTD/MND was driven by an increased proportion of expressing cells, with the strongest effects observed in L2/3 IT, L5 ET, and L5/6 IT populations. Boxplots represent distributions across individuals.

**Extended Data Fig. 7.**
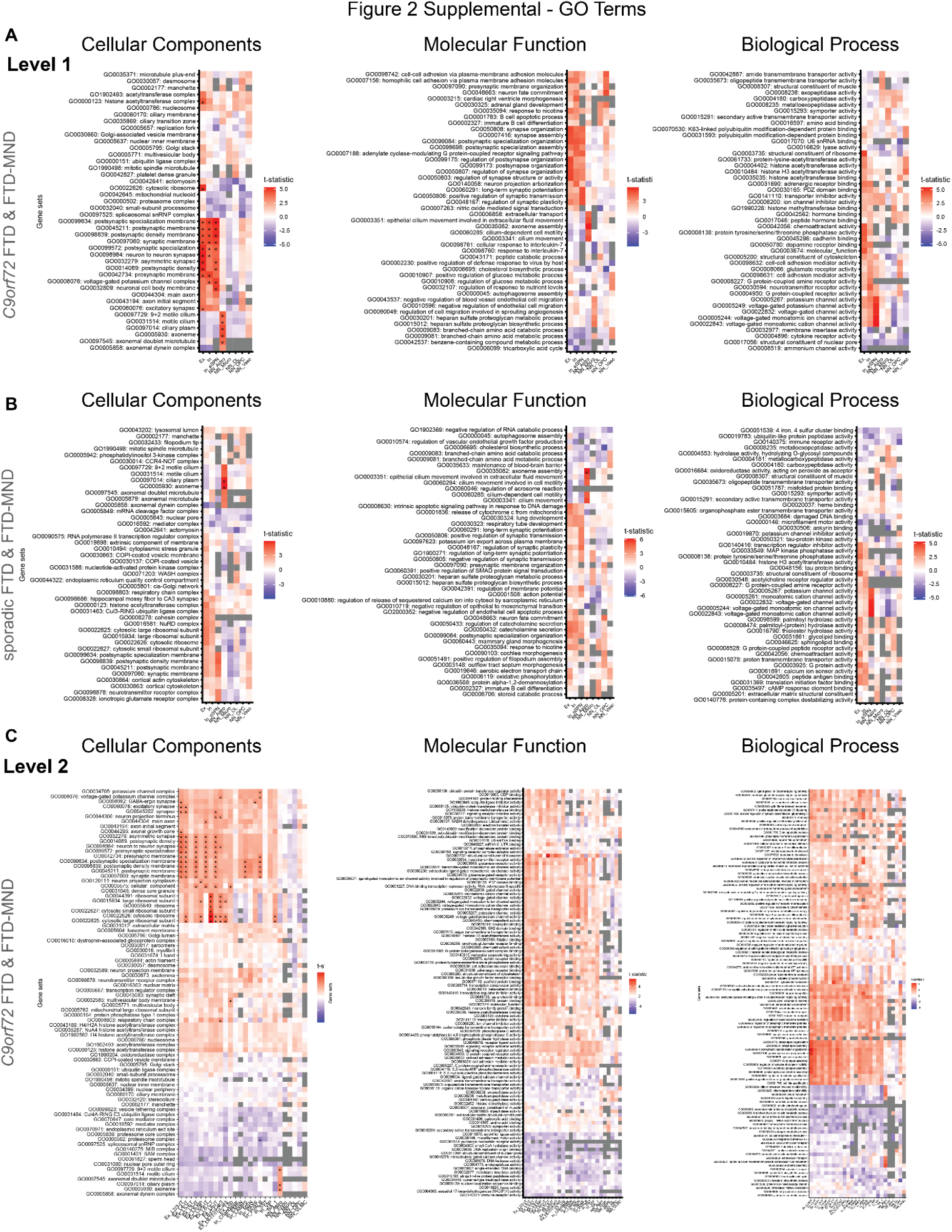
Convergent pathway remodeling across FI excitatory neuronal populations in *C9orf72* and sporadic FTD & FTD-MND. **A:** Gene ontology (GO) enrichment analysis across broad cell classes (Level 1 taxonomy) in *C9orf72* FTD/FTD-MND. Heatmaps show enriched cellular component, molecular function, and biological process gene programs across major FI cell populations. Excitatory neurons showed enrichment of synaptic, dendritic, axonal, and ion channel-related pathways, whereas astrocytes showed enrichment of cilia-, axoneme-, and microtubule-related programs. Colors indicate enrichment t-statistics. **B:** GO enrichment analysis across broad cell classes (Level 1 taxonomy) in sporadic FTD/FTD-MND. Pathway alterations were less significant, but broadly concordant with those observed in *C9orf72* disease, including enrichment of synaptic and neuronal structure-related programs in excitatory neurons and cilia-related programs in astrocytes. Colors indicate enrichment t-statistics. **C:** GO enrichment analysis across excitatory neuron subclasses (Level 2 taxonomy) in *C9orf72* FTD/FTD-MND. Excitatory neuron subclasses exhibited broad upregulation of synaptic, dendritic, axonal, ribosomal, and signaling-related gene programs across disease. Colors indicate enrichment t-statistics.

**Extended Data Fig. 8.**
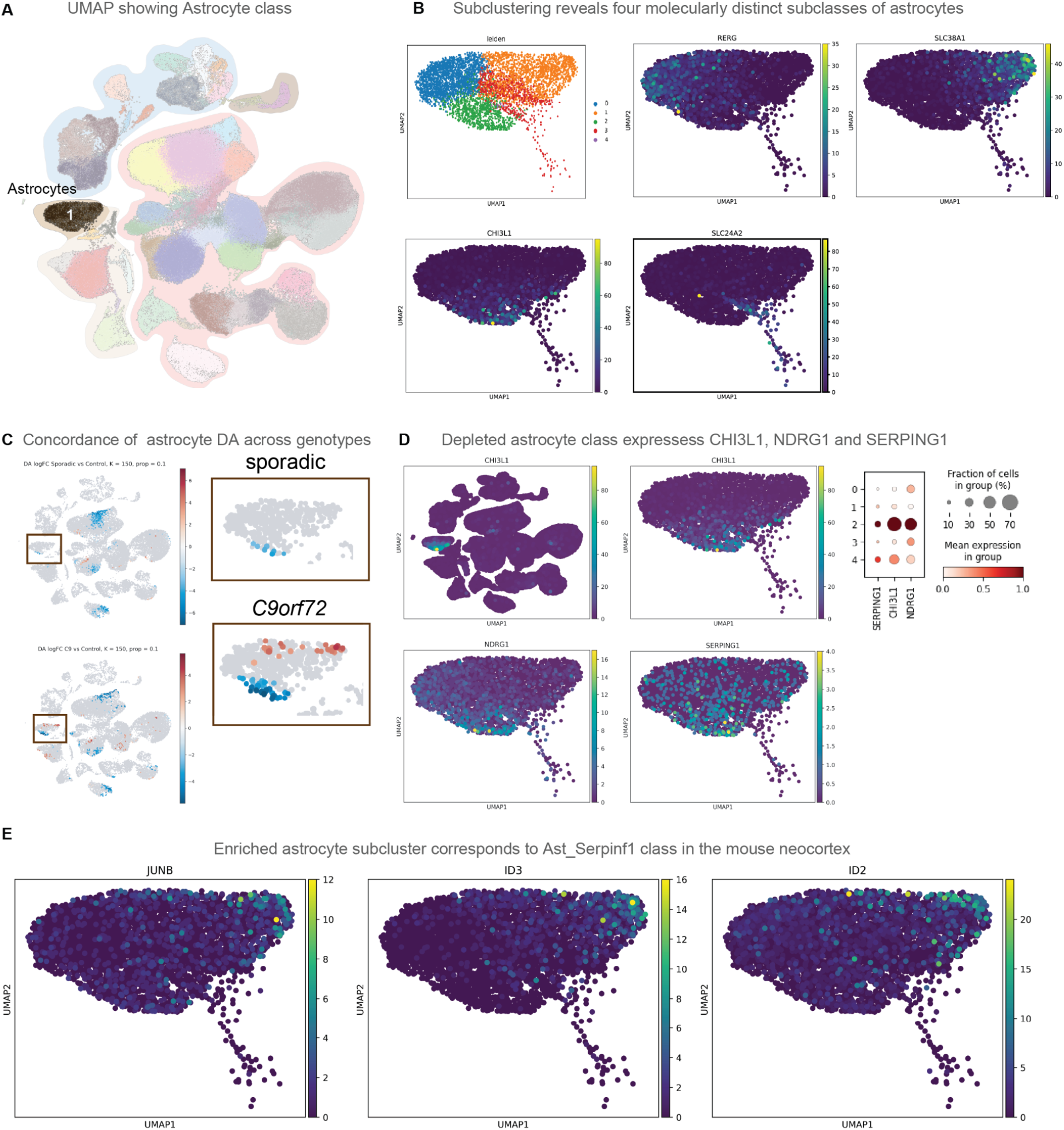
Reactive astrocyte subclasses are differentially represented in FTD & FTD-MND. **A:** UMAP highlighting the astrocyte population within our integrated FI snRNA-seq dataset. **B:** Subclustering analysis identifies four molecularly distinct astrocyte subclasses demarcated by expression of RERG, SLC38A1, CHI3L1, and SLC24A2. **C:** Differential abundance analysis reveals concordant depletion of astrocyte subcluster 2 across *C9orf72* and sporadic FTD/FTD-MND. **D:** Astrocyte subcluster 2 highly expresses reactive astrocyte-associated markers, including CHI3L1, NDRG1, and SERPING1. **E:** Astrocyte subcluster 1, which is relatively enriched in disease, highly expresses JUNB, ID2, and ID3, corresponding to the previously described Ast_Serpinf1 astrocyte class in the mouse neocortex.

**Extended Data Fig. 9.**
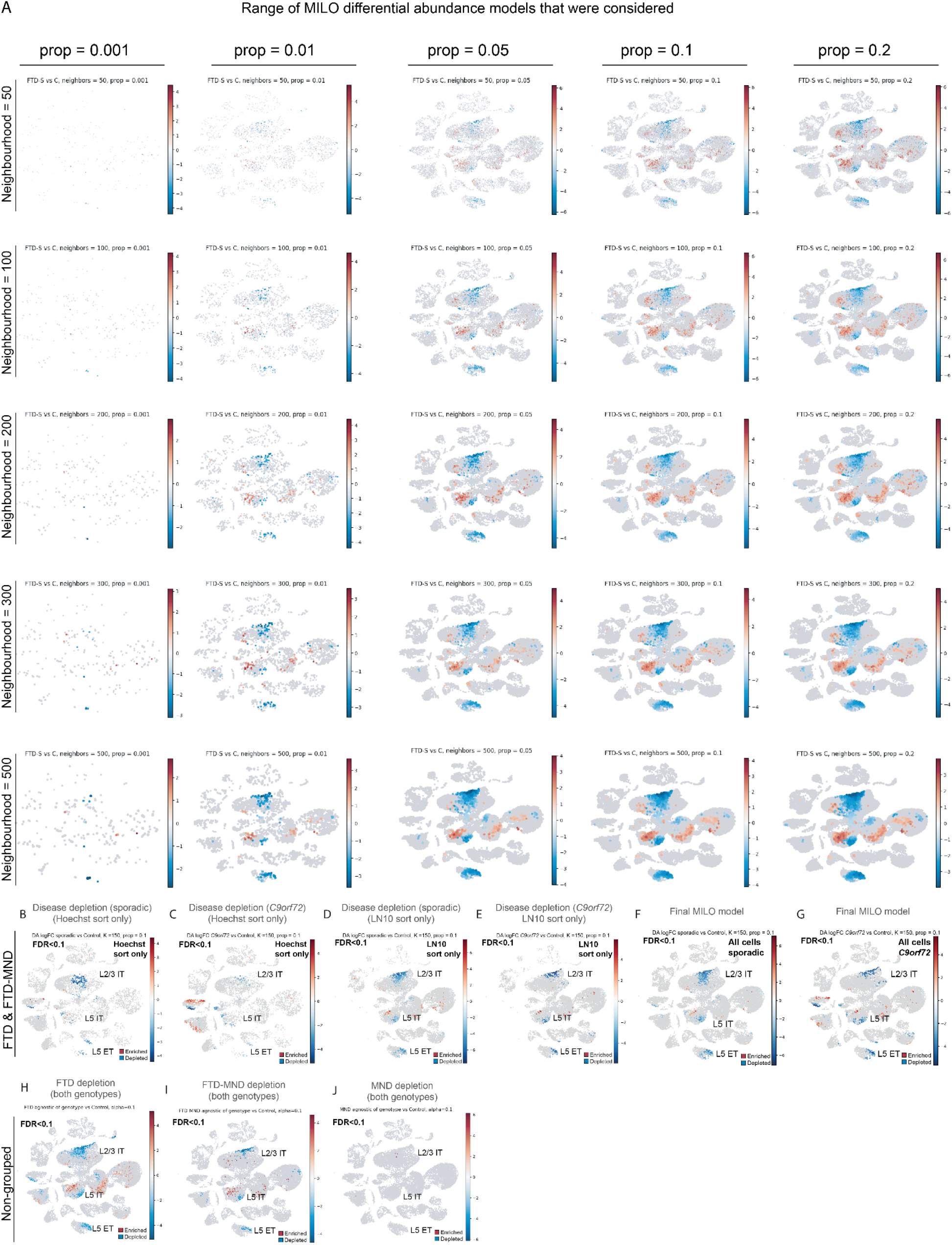
MILO differential abundance signatures are robust across model parameters and sorting conditions. **A:** MILO differential abundance (DA) analyses performed across a range of neighborhood sizes and minimum neighborhood proportion thresholds. Consistent DA patterns were observed across a broad parameter space, including depletion and enrichment signatures among excitatory neuron populations. **B-E:** MILO DA analyses of grouped FTD/FTD-MND cases performed separately on unenriched Hoechst+ nuclei (B–C) and LN10-enriched nuclei (D–E) for sporadic (B,D) and *C9orf72* (C,E) disease groups. DA patterns were broadly concordant across sorting and genotype conditions. **F-G:** Final MILO DA models for sporadic FTD/FTD-MND (F) and *C9orf72* FTD/FTD-MND (G) using all nuclei and a neighborhood size of 150 with a minimum neighborhood proportion threshold of 0.1. Significant depletion (FDR < 0.1) of L2/3 IT, L5 IT, and L5 ET populations was observed across disease groups. **H-I:** MILO DA analyses performed separately on clinically defined FTD (H) and FTD-MND (I) groups revealed depletion patterns broadly consistent with the grouped FTD/FTD-MND analyses. **J:** MILO DA analysis of clinically pure MND cases identified minimal differential abundance changes in the FI, consistent with the preferential vulnerability of the primary motor cortex in MND.

**Extended Data Fig. 10.**
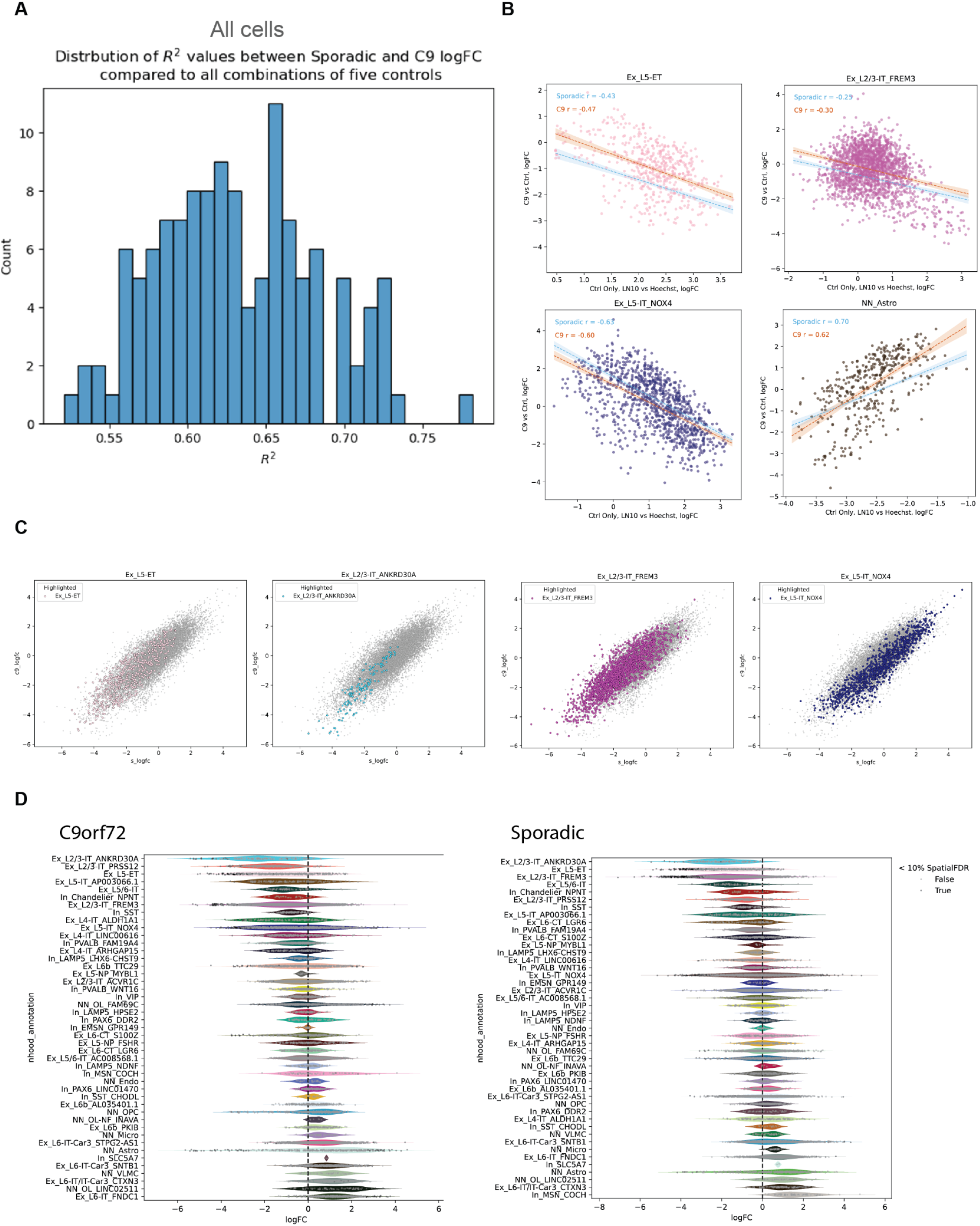
Shared vulnerability of large excitatory neuron populations across *C9orf72* and sporadic FTD/MND is only partially explained by nuclear size. **A:** Distribution of Pearson correlation (R-squared) values comparing neighborhood-level differential abundance logFCs between *C9orf72* and sporadic FTD & FTD-MND relative to all combinations of five control samples, demonstrating robust concordance of disease-associated depletion patterns across genotypes. **B:** Relationship between LN10 sorting enrichment score, a proxy for neuronal size, and disease-associated differential abundance across representative excitatory neuron populations. L5 ET, L2/3 IT FREM3, and L5 IT NOX4 populations showed modest negative correlations between size enrichment and disease-associated abundance. **C:** Neighborhood-level disease-associated differential abundance logFC values were highly concordant across *C9orf72* and sporadic FTD & FTD-MND within vulnerable excitatory neuron populations, including L5 ET, L2/3 IT ANKRD30A, L2/3 IT FREM3, and L5 IT NOX4. **D:** Distribution of neighborhood-level differential abundance logFC values across FI cell populations in *C9orf72* and sporadic FTD & FTD-MND. Vulnerable excitatory neuron populations, including L2/3 IT, L5 IT, and L5 ET subclasses, showed the strongest depletion in disease.

**Extended Data Fig. 11.**
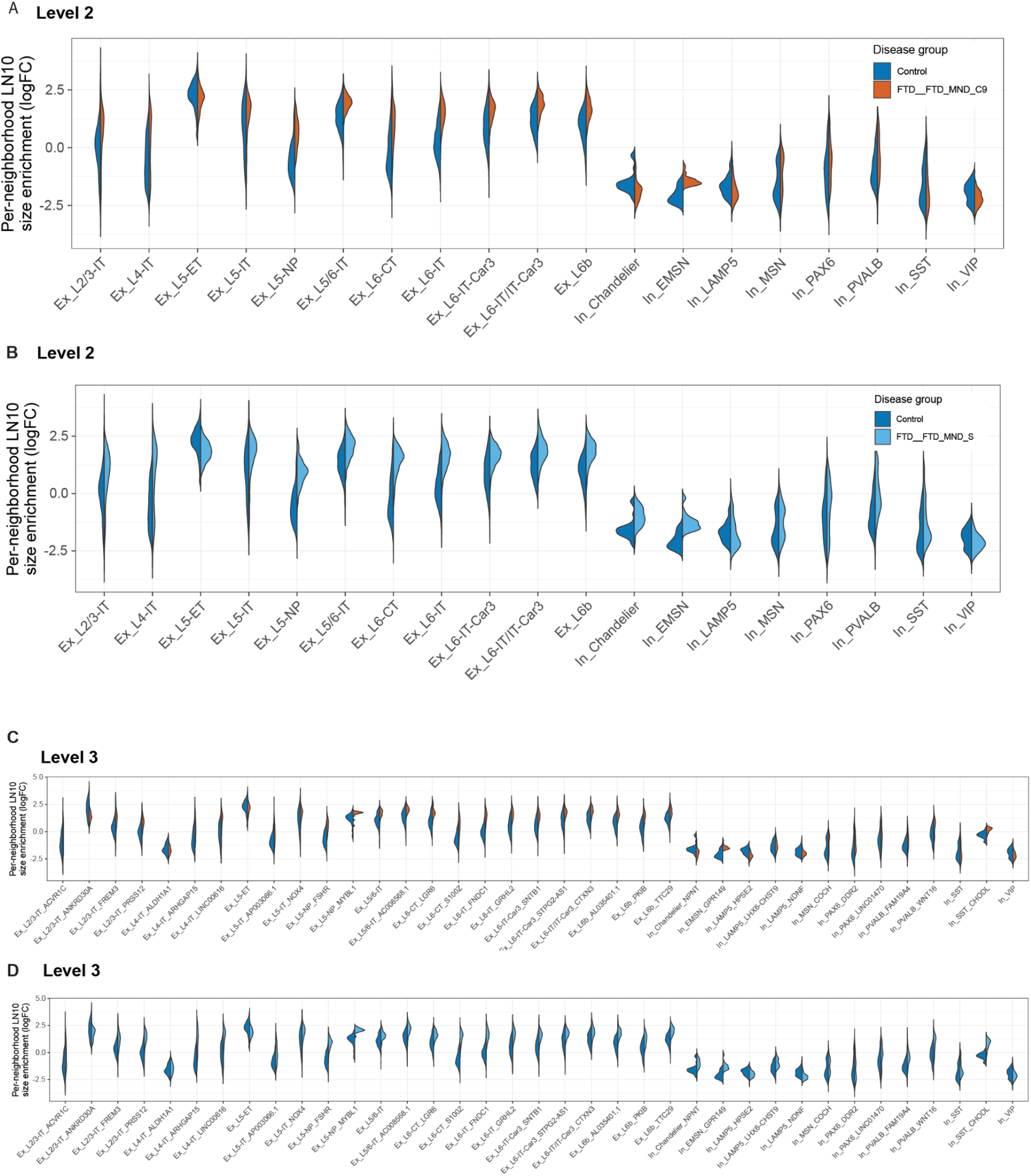
Disease-associated neuronal enlargement extends broadly across excitatory neuron populations. **A:** Neighborhood-level LN10 enrichment scores, used as a proxy for neuronal size, across Level 2 excitatory and inhibitory neuron populations in control and *C9orf72* FTD & FTD-MND samples. Broad excitatory neuron populations exhibited increased LN10 enrichment in disease relative to controls, whereas inhibitory neuron populations showed comparatively limited changes. **B:** Neighborhood-level LN10 enrichment scores across Level 2 excitatory and inhibitory neuron populations in control and sporadic FTD & FTD-MND samples. Broad excitatory neuron populations showed increased LN10 enrichment in disease. **C:** Neighborhood-level LN10 enrichment scores across Level 3 neuronal and non-neuronal populations in control and *C9orf72* FTD & FTD-MND samples. Disease-associated size enrichment extended across multiple excitatory neuron subclasses and cortical layers. **D:** Neighborhood-level LN10 enrichment scores across Level 3 neuronal and non-neuronal populations in control and sporadic FTD & FTD-MND samples. Broad disease-associated size enrichment was observed across excitatory neuron subclasses despite selective vulnerability of only specific populations.

**Extended Data Fig. 12.**
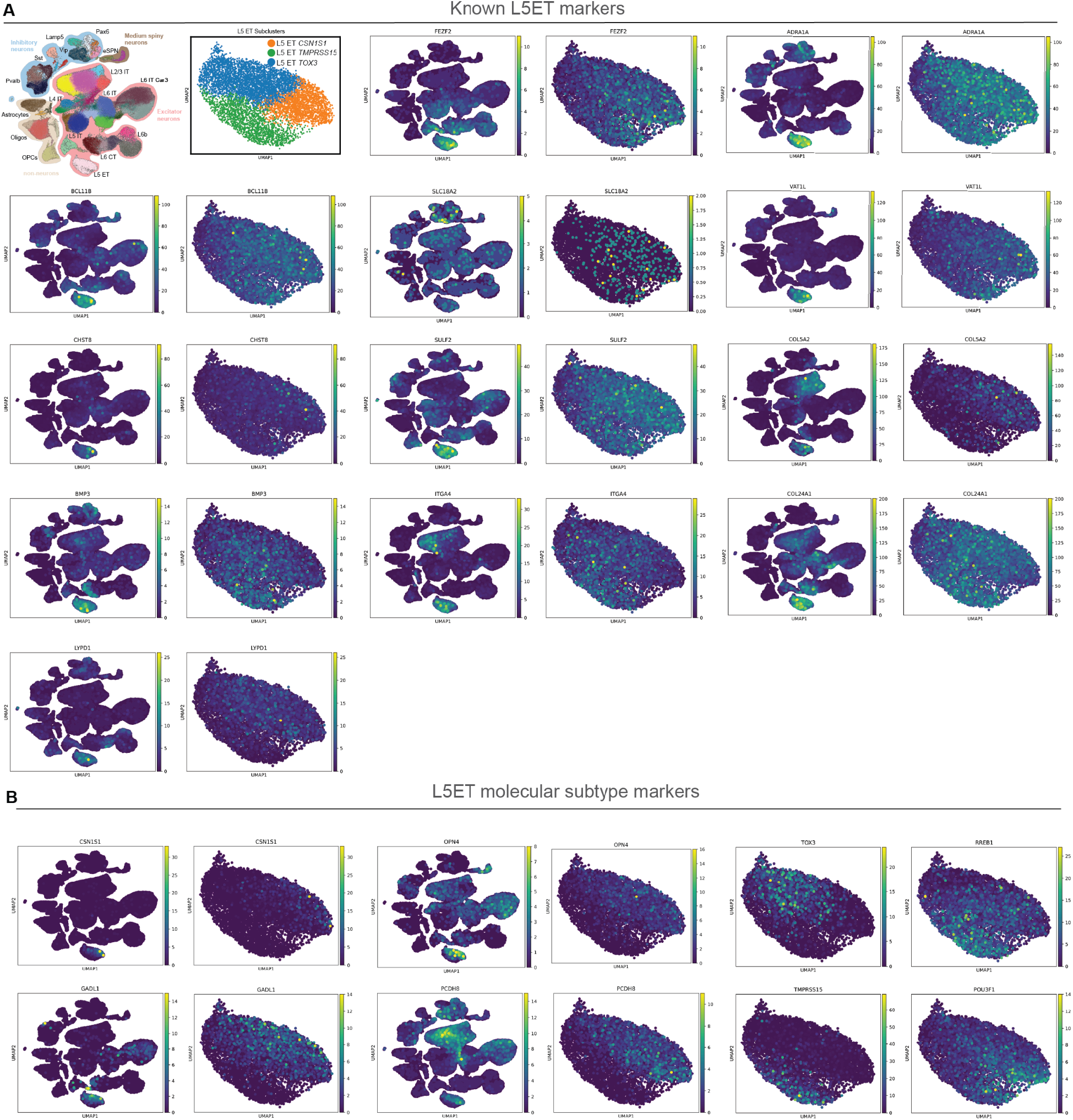
Novel subtype markers distinguish molecularly distinct L5 ET populations. **A:** Expression of previously reported VEN/L5 ET-associated marker genes across the full FI dataset and within isolated L5 ET neurons. Most published VEN markers, including *FEZF2*, *ADRA1A*, *VAT1L*, *CHST8*, *SULF2*, *BMP3*, *ITGA4*, *COL5A2*, *COL24A1*, and *LYPD1*, were broadly expressed across L5 ET subclasses and did not distinguish molecularly defined L5 ET subtypes. **B:** Expression of candidate subtype-specific markers across L5 ET subclasses. L5 ET *CSN1S1* cells were characterized by expression of *CSN1S1*, *GADL1*, *OPN4*, and PCDH8. L5 ET *TOX3* cells were characterized by expression of *TOX3*, *GADL1*, *OPN4*, and PCDH8. L5 ET *TMPRSS15* was characterized by expression of *TMPRSS15*.

**Extended Data Fig. 13.**
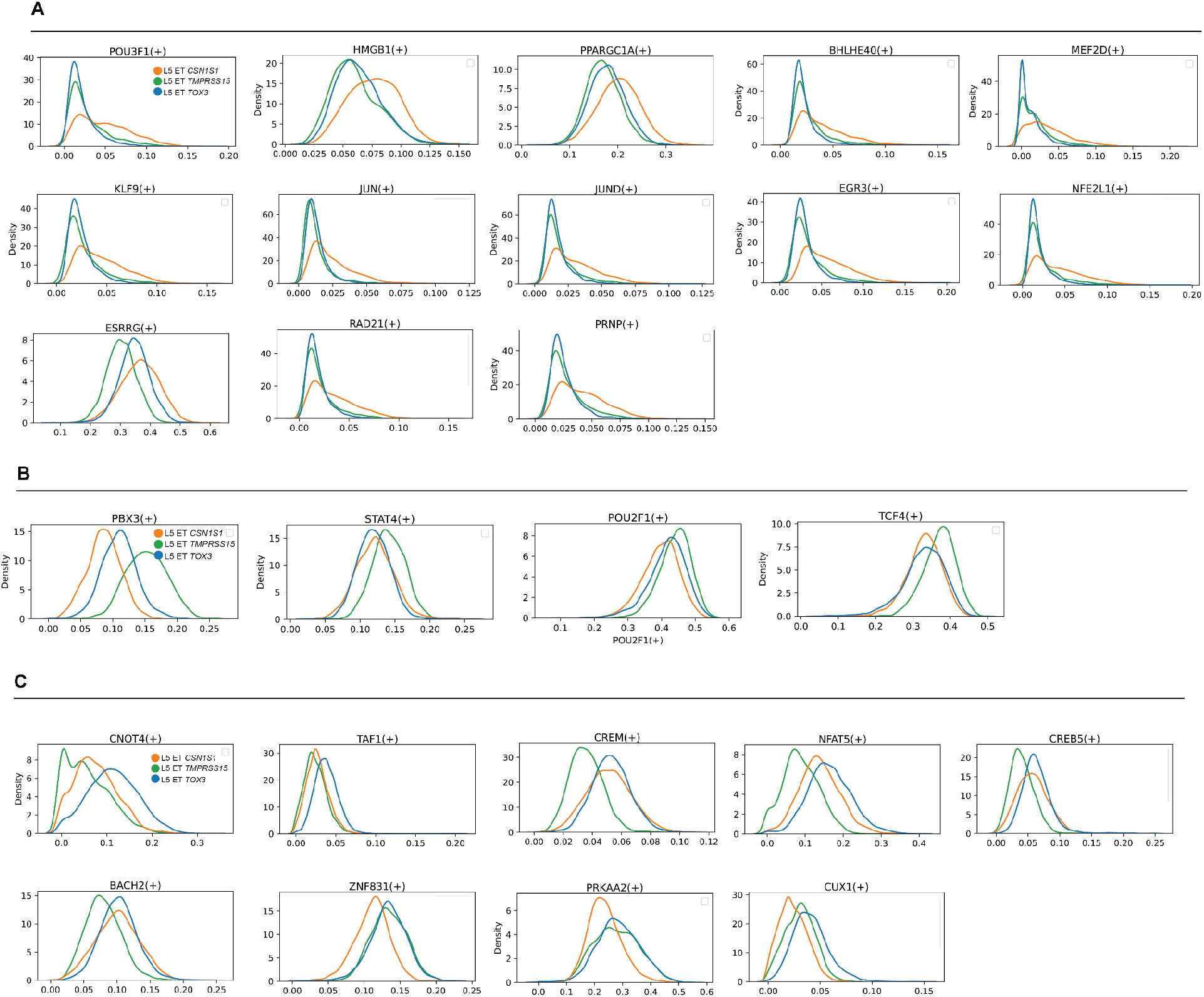
Subtype-specific regulon enrichment across L5 ET neuron populations. **A:** Gene regulatory network analysis identifies distinct regulon enrichment patterns across L5 ET subclasses. L5 ET CSN1S1 neurons were enriched for regulons associated with *POU3F1*, *HMGB1*, *BHLHE40*, *MEF2D*, *KLF9*, *JUN*, metabolic regulators *PPARGC1A* and *ESRRG*, early response genes *JUND* and *EGR3*, and oxidative stress response gene *NFE2L1*. **B:** L5 ET TMPRSS15 was distinguished by *PBX3*, *STAT4*, *POU2F1*, and *TCF4*, consistent with transcriptional programs involved in neuronal identity maintenance and responsiveness to extracellular signaling pathways. **C:** L5 ET TOX3 was distinguished by *CNOT4* and *TAF1*, factors involved in mRNA turnover and transcriptional initiation, respectively, and shared many marker genes with L5 ET CSN1S1 that were less highly expressed in the L5 ET TMPRSS15 class, such as *CREM*, *NFAT5*, *CREB5* and *BACH2*, indicating high homology between these two classes and the relatively distinct transcriptional nature of the TMPRSS15 class

**Extended Data Fig. 14.**
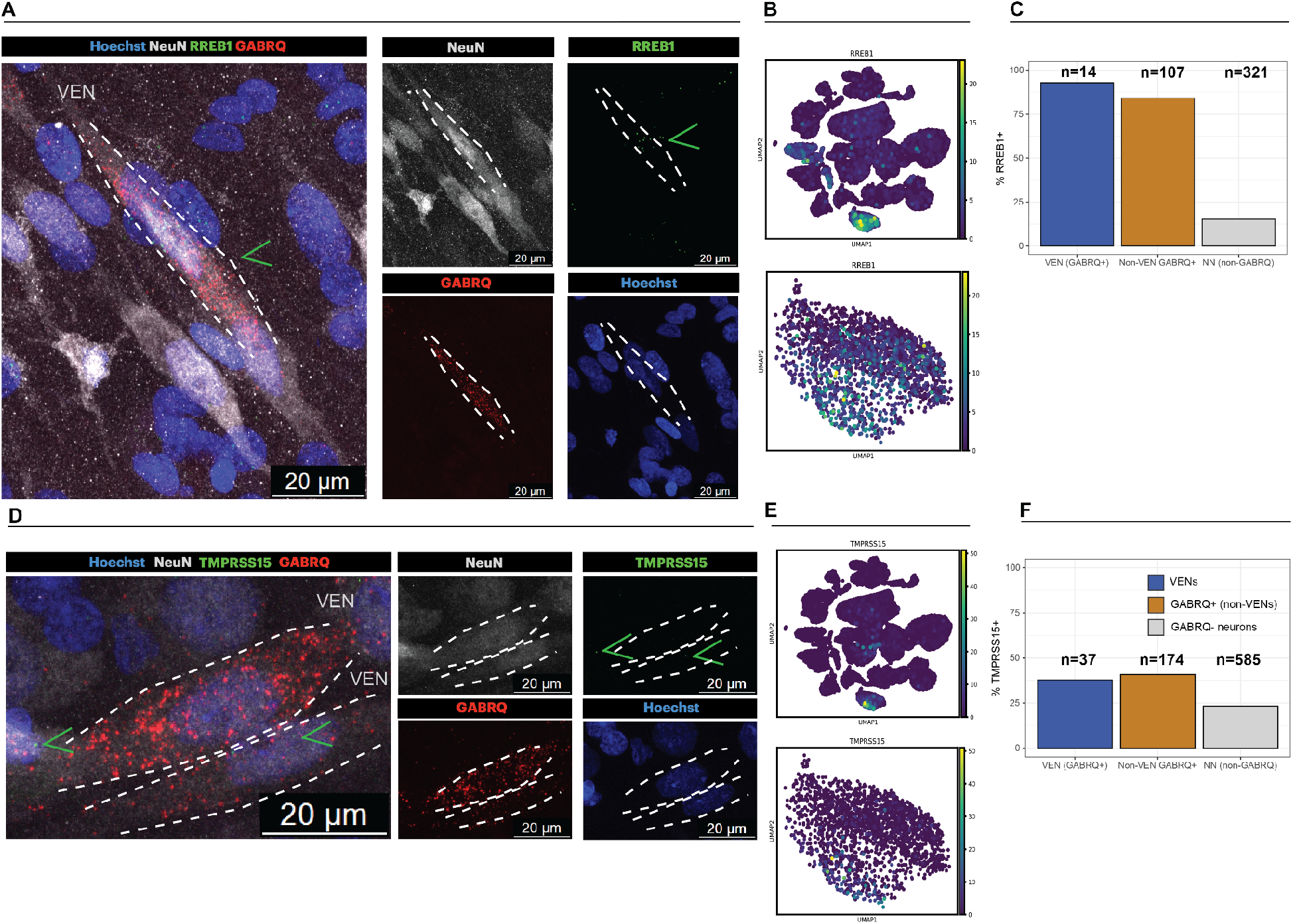
RREB1 and TMPRSS15 expression do not separate morphologically defined VENs from non-VEN GABRQ+ neurons. **A:** Multiplex fluorescent RNAscope showing co-expression of RREB1 in GABRQ+ neurons within a morphologically defined VEN in the FI. Representative images show Hoechst, NeuN, RREB1, and GABRQ. Note that some non-neuronal cells also express RREB1. These are likely glial cells (astrocytes) since transcriptional expression of RREB1 is also found in subsets of glial cells (panel B). Scale bars, 20 μm. **B:** UMAPs showing RREB1 expression is highly expressed in the L5 ET class with shared expression across the three L5 ET sub clusters (L5 ET *TMPRSS15*, L5 ET *TOX3* and L5 ET *CSN1S1*). Note that subsets of glial cells (e.g. astrocytes) also express RREB1. **C:** RREB1 is expressed in 93% of VENs (13/14) and 84% of non-VEN GABRQ+ neurons (90/107), but does not differentiate the two populations. **D:** Multiplex fluorescent RNAscope showing co-expression of TMPRSS15 and GABRQ in one of two shown VENs within the FI. On the left is an example of a non-VEN GABRQ+ neuron that expresses TMPRSS15. Representative images show Hoechst, NeuN, TMPRSS15, and GABRQ. Arrowheads indicate a TMPRSS15-positive VEN and non-VEN GABRQ+ neuron. Scale bars, 20 μm. **E:** UMAPs showing that TMPRSS15 expression is mostly unique to the L5 ET cluster, and is most highly expressed in the L5 ET *TMPRSS15* cluster. **F:** TMPRSS15 was expressed in 37.8% of VENs (14/37), 40.8% of non-VEN GABRQ+ cells (71/174) and 23.2% of neighboring neurons (136/585).

**Extended Data Fig. 15.**
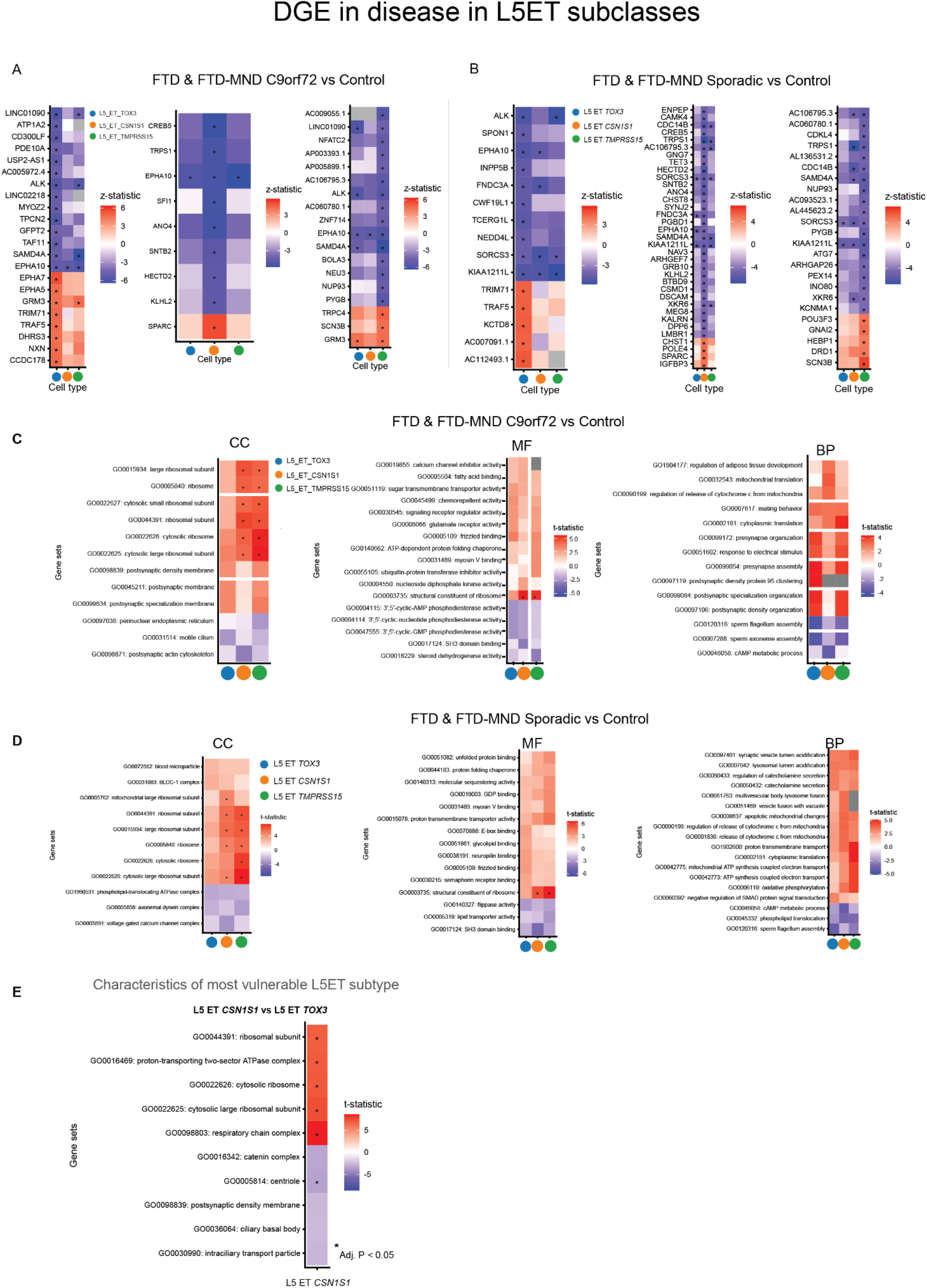
Oxidative phosphorylation programs distinguish the most vulnerable L5 ET subtype (L5 ET *CSN1S1*) **A:** Differential gene expression analysis of L5 ET subclasses in *C9orf72* FTD & FTD-MND relative to controls. Heatmaps show significantly altered genes across L5 ET *TOX3*, L5 ET *CSN1S1*, and L5 ET *TMPRSS15* populations. Colors indicate z-statistics. Orange represents L5 ET *CSN1S1*. Blue represents L5 ET *TOX3*. Green represents L5 ET *TMPRSS15*. **B:** Differential gene expression analysis of L5 ET subclasses in sporadic FTD & FTD-MND relative to controls. Disease-associated transcriptional changes were broadly shared across L5 ET subclasses and genotypes. Colors indicate z-statistics. **C:** Gene ontology enrichment analysis of disease-associated transcriptional changes across L5 ET subclasses in *C9orf72* FTD & FTD-MND. Cellular component (CC), molecular function (MF), and biological process (BP) enrichments are shown. Colors indicate enrichment t-statistics. **D:** Gene ontology enrichment analysis of disease-associated transcriptional changes across L5 ET subclasses in sporadic FTD & FTD-MND. Disease-associated pathway alterations were broadly shared across L5 ET subclasses and genotypes. Colors indicate enrichment t-statistics. **E:** Gene ontology enrichment analysis comparing control L5 ET *CSN1S1* and L5 ET *TOX3* neurons. The more vulnerable L5 ET *CSN1S1* subtype was enriched for oxidative phosphorylation, respiratory chain, ATP synthesis, and ribosomal gene programs, whereas L5 ET *TOX3* neurons showed relative enrichment of centriole- and cilia-associated pathways. Asterisks indicate adjusted P < 0.05.

## Supplementary Tables

**Supplementary Tables:** https://ucsf.box.com/s/04230hrm0r9f6wqnuekgapm1q5k5iutc

**Supplementary Table 1**: Case cohort (1a), Cell Type information (1b), Cell Type Marker Genes (1c-e)

**Supplementary Table 2**: Cell type-specific differential expression level 1 (2a), DGE level 2-3 (2b), GO terms (2c)

**Supplementary Table 3**: MiloDA (3a), GE-DA (3b)

**Supplementary Table 4**: Cell Type Marker Genes L5 ET subtypes (4a-c), quantifications RNAscope (4d-e)

## Methods

### Human post-mortem tissue collection

Frozen post-mortem frontoinsular cortex (FI) tissue was obtained from the UCSF Neurodegenerative Disease Brain Bank. Patients or their surrogates provided informed consent for brain donation and genetic testing prior to autopsy, in keeping with the guidelines put forth in the Declaration of Helsinki. Neuropathological diagnoses were made following consensus diagnostic criteria (Mackenzie et al., 2010; Montine et al., 2011) using previously described histological and immunohistochemical methods (Kim et al., 2012; Tartaglia et el., 2010). Healthy control tissues were obtained from individuals without dementia who had minimal age-related neurodegenerative changes. Genetic screening was performed for common and rare FTD-related genes as previously described (Ramos et al., 2020). Cases were identified based on neuropathological diagnosis and genetic screening.

The cohort comprised 40 individuals, including patients with C9orf72-associated FTLD-TDP Type B or Type U/MND-TDP pathology (n = 16), sporadic FTLD TDP Type B/MND-TDP pathology (n = 15), and neuropathologically normal controls (n = 9) (Fig.1 a & Suppl. Table 1a). To promote pathological and clinical homogeneity, enrollment was restricted to individuals spanning the bvFTD–MND clinical spectrum. Cases were selected to represent a range of FI involvement, from clinically pure MND with minimal extramotor TDP-43 pathology to advanced bvFTD with extensive FI degeneration and TDP-43 burden. Samples were selected to minimize technical confounds, including comparable post-mortem intervals (PMI) and RNA integrity numbers (RIN), and were processed into a single pooled batch during nuclei isolation and fluorescence activated nuclei sorting to reduce batch effects.

### Nuclei isolation of fresh frozen frontoinsular cortex samples

Nuclei were isolated from frozen FI tissue using a sucrose-gradient ultracentrifugation protocol. All procedures were performed under RNase-free conditions and on ice. Briefly, sectioned or finely chopped frozen tissue was homogenized on ice in 5mL Lysis buffer containing sucrose (0.32M), CaCl_2_ (3mM), Mg(Ac)_2_ (3mM), EDTA (0.1mM), Tris-HCl (pH 8) (10mM), DTT (1mM), Triton X-100 (0.1%) and DEPC-water using a glass Dounce homogenizer (Thomas Scientific; Catalog # 3431D76; Size A). Homogenates were transferred to thick-wall ultracentrifuge tubes (Beckman Coultier; Catalog # 355631) and carefully layered over 9mL Sucrose solution containing sucrose (1.8M), Mg(Ac)_2_ (3mM), DTT (1mM), Tris-HCl (10mM) and DEPC-water to generate distinct phases. Samples were balanced by weight and centrifuged in a Beckman SW28 swing-bucket rotor at 24,400 r.p.m. (107,163.6g) for 1.5 h at 4 °C. Following ultracentrifugation, supernatants was carefully removed and nuclei at the bottom of the ultracentrifuge tubes were resuspended in DEPC-treated PBS on ice prior to filtration through 37 μm filters. Nuclei were counted using a Countess cell counter, and aliquots were reserved for nuclear morphology assessment.

### Fluorescence-activated nuclei sorting (FANS) of isolated nuclei

Isolated nuclei were stained and sorted using a fluorescence-activated nuclei sorting (FANS) workflow adapted for adult human post-mortem brain tissue. Following nuclei isolation, nuclei concentrations were quantified using a Countess cell counter. Nuclei were permeabilized and stained in buffer containing human FcX blocker (1:100; BioLegend, catalog #156604), ultrapure BSA (1%; Thermo AM2618), Tween-20 (0.2%), RNAse inhibitor (0.7 units μL/uL; Millipore Sigma, catalog #3335399001) and PBS. The staining buffer was adapted from a previous publication ^76^, with dextran sulfate omitted to reduce nuclei aggregation. Nuclei were incubated in staining buffer at 4 °C for 10 min, followed by staining with Hoechst and NeuN-Alexa647 antibody (1:1,000; Abcam, catalog #ab190565) for 30 min at 4 °C in the dark on a rotating platform. NeuN-only, Hoechst-only and unstained controls were included. Nuclei were subsequently washed three times in PBS containing 1% BSA and RNase inhibitor by centrifugation at 500g for 5 min at 4 °C.

FANS was performed using a BD Fusion cell sorter equipped with a 130 μm nozzle to preserve nuclear integrity. Hoechst fluorescence was detected using UV/violet excitation and NeuN-Alexa647 using APC excitation. To enrich for the large L5 ET neuron class while retaining broad FI cellular diversity, nuclei were sorted into two complementary populations: a non-enriched Hoechst+ condition (Hoechst) and an LN10-enriched condition (LN10) consisting of approximately the largest 10% of NeuN+ neuronal nuclei based on Forward Scatter Area (FSC-A) signal. Debris and aggregates were excluded using SSC-A, FSC-A and FSC-W gating. Putative single nuclei were identified using FSC-W and FSC-A gating, and further refined by selecting a discrete Hoechst+ population. Large neuronal nuclei were next enriched by gating the ∼10% largest neuronal nuclei according to the FSC-A signal. Following sorting, nuclei were centrifuged at 500g for 5 min at 4 °C, resuspended in PBS/BSA buffer, quantified using a Countess cell counter and adjusted to target concentrations for downstream single-nucleus RNA sequencing workflows.

### Single-nucleus RNA sequencing

Single-nucleus RNA-seq (snRNA-seq) libraries were generated using the Chromium Single Cell 3′ Gene Expression v3.1 HT platform (10x Genomics, Catalog #1000370) according to the manufacturer’s instructions (CG000417, Rev D). Libraries were sequenced on Illumina instruments. Using contemporary best-practice guidelines for single-cell analysis (^77^), quality-control filtering was performed in Scanpy to remove low-quality nuclei and low-abundance genes. Genes detected in fewer than five nuclei were excluded. Nuclei with fewer than 1,000 total UMI counts, greater than 150,000 total UMI counts, or greater than 5% mitochondrial transcript content were also removed. Putative doublets and ambient RNA contamination were subsequently identified using genotype-based demultiplexing approaches (see below). Following quality control, 224,793 nuclei were retained for downstream analyses, including 85,635 nuclei from the Hoechst+ condition and 139,158 nuclei from the LN10-enriched condition.

### Extraction and genotyping of genomic DNA

Genomic DNA was extracted from snap-frozen adult human brain tissue using the PureLink Genomic DNA Mini Kit (Thermo Fisher Scientific, catalog #K182001) according to the manufacturer’s protocol with modifications optimized for lipid-rich adult human brain tissue. Frozen tissue was finely dissected prior to extraction, and approximately 10 mg of tissue was used per reaction. To improve digestion and DNA yield, 30 μL of Proteinase K was used instead of the manufacturer-recommended 20 μL, and samples were digested at 55 °C on a shaking thermomixer (600 r.p.m.) for approximately 4 h with intermittent pipetting and brief vortexing every 30 min to facilitate homogenization. Binding steps were performed only after homogenates appeared fully uniform without visible tissue fragments. Following the second wash step, columns were centrifuged for an additional 1.5 min at maximum speed to remove residual wash buffer. To maximize DNA recovery, the initial eluate was reapplied onto the spin column and centrifuged again for 1.5 min at maximum speed. These modifications improved DNA yield and quality from frozen adult human brain tissue relative to the standard manufacturer protocol. DNA samples were subsequently genotyped using the Infinium Global Screening Array platform (Illumina, catalog #20030770) according to the manufacturer’s protocol for Infinium HTS assays.

### Genetic demultiplexing, ambient RNA estimation, and doublet detection

To assign nuclei to individual-of-origin and estimate ambient RNA contamination, we leveraged naturally occurring genetic variation, derived from the Global Screening Array Platform (see above), using CellBouncer^31^. Specifically, the demux_vcf workflow was used to probabilistically assign nuclei to donors on the basis of expressed single-nucleotide polymorphisms derived from matched genotype VCF files, allowing us to demultiplex the 40 individuals in our pooled data with high confidence. We performed donor demultiplexing, identified putative doublets, and estimated ambient RNA contamination across sequencing lanes following standard procedures. All individuals were successfully recovered following demultiplexing and doublets were removed.

### Data integration and clustering

RNA sequencing data was processed and analyzed using Scanpy ^78^. To remove separation of the same cell types caused by confounding variables we integrated the dataset using scVI ^32^. We selected the top 5000 highly variable features, and controlled for Clinical Grouping x Gene Status, Individual and Sex as categorical covariates, and age at death and post-mortem interval as continuous covariates. For the Clinical Grouping x Gene Status category, we grouped clinical bvFTD and bvFTD-MND together given their shared involvement of FI and split them into *C9orf72* and sporadic groups. The default scVI settings were used for the neural network size (n_hidden: 128, n_latent: 10, n_layers: 1, dropout_rate: 0.1, dispersion: gene, gene_likelihood: zinb, latent_distribution: normal). We used the latent space created by scVI to build a neighborhood graph and applied unifold manifold approximation and projection (UMAP) to reduce dimensionality and visualize the cell position in two-dimensional space. Raw gene expression counts per cell were first normalized to the median of total counts across cells (scanpy.pp.normalize_total()), then logarithmically transformed to ease visualization and downstream analysis (scanpy.pp.log1p()).

Graph-based clustering was performed using the Leiden algorithm with resolution set to 2. To annotate clusters we inspected the first 500 marker genes derived with the logistic regression algorithm, and merged clusters which lacked unique marker genes. Clusters were retained on the basis of transcriptionally distinct marker gene expression profiles, resulting in 47 molecularly defined cell populations (Suppl. Table 1b: cell level 3). Cell type annotations were additionally (Suppl. Table 1b: cell level 2) assigned through reference mapping with MapMyCells (RRID:SCR_024672) against the Allen Institute Whole Human Brain taxonomy and the Human MTG SEA-AD taxonomy ^25,26^. Hierarchical annotations were further organized into multiple taxonomic resolutions (cell levels 1–3; Suppl. Table 1b). To identify molecular subtypes in L5 ET cells and in astrocytes, we subclustered with resolution parameters 0.15 and 0.2, respectively, and annotated marker genes based on logistic regression.

### Differential gene expression analysis

To analyze differential expression patterns, we utilized precision-weighted linear models implemented in Dreamlet ^40^. Prior to this analysis, we converted the h5ad scanpy object to a RDS SingleCellExperiment object using zellkonverter. To mitigate sparsity and pseudoreplication inherent to single-cell datasets, raw gene expression counts were aggregated into pseudobulk profiles using dreamlet::aggregateToPseudoBulk() within transcriptionally defined populations across cell levels 1, 2 and 3 (Suppl. Table 1b) for each unique sample defined as the combination of Individual, Sex and Clinical Grouping (Clinical Diagnosis × Gene Status). Prior to modeling, age at death and post-mortem interval (PMI) were z-score scaled by subtracting the mean and dividing by the standard deviation. Processed assays were generated using dreamlet::processAssays() with min.count = 5 and min.prop = 0.2, retaining genes expressed in at least 20% of samples within each assay (cell type). Differential expression was modeled using the following fixed-effects formula: *∼0 + ClinGene_DEG_group + Sex + age_scaled + pmi_scaled*. Contrasts were computed separately for sporadic FTD&FTD-MND, *C9orf72* FTD&FTD-MND, sporadic MND and *C9orf72* MND relative to controls, as well as between sporadic FTD&FTD-MND relative to *C9orf72* FTD&FTD-MND, and sporadic MND relative to *C9orf72* MND. Differential expression analyses were performed at both coarse and fine taxonomic resolutions (cell levels 1 and 3; Suppl. Table 2a-b). Genes were considered differentially expressed following false discovery rate (FDR) correction (adjusted P < 0.05) performed within cell types.

### Gene ontology and pathway enrichment analyses

Gene ontology and pathway enrichment analyses were performed using the full distribution of gene-level differential expression statistics generated by Dreamlet^40^ (see Differential gene expression analysis). Rather than thresholding genes into significant and non-significant categories prior to enrichment testing, we instead performed gene set analysis using the Zenith:gsa() function inside Dreamlet ^40^, a rotation-based gene set testing framework built on limma::camera that evaluates enrichment using the full distribution of differential expression statistics while accounting for inter-gene correlation within pathways. Enrichment analyses were conducted separately for major cell classes and transcriptionally defined subtypes to identify altered biological programs associated with disease, including synaptic, dendritic, axonal, ciliary, inflammatory, and oxidative phosphorylation pathways. Gene set enrichment analysis (GSEA) was performed using fgsea ^79^ implemented through ClusterProfiler ^80^, and terms were hierarchically collapsed using rrvgo^81^.

### Differential abundance analysis

Differential abundance (DA) analysis was performed using Milo^30^ implemented through pertpy. Following Milo parameter recommendations, neighborhood graphs were constructed in the scVI latent space using a k-nearest neighbor graph with k = 150 nearest neighbors and prop = 0.1, corresponding to sampling 10% of cells as neighborhood index cells, resulting in 16,342 neighborhoods. To assess relative neuronal size enrichment, neighborhood counts were aggregated using count_nhoods(sample_col=“batch”), and DA testing was performed comparing LN10 and Hoechst sorting conditions using the model design=∼condition with the contrast conditionLN10-conditionHoechst. Neighborhood-level LN10 enrichment log fold-change values were subsequently used as a proxy for relative neuronal size.

To assess disease-associated DA, neighborhood counts were aggregated at the individual level using count_nhoods(sample_col=“Individual”). Neighborhoods were annotated using annotate_nhoods() with the cell_level_3 taxonomy, and DA testing was performed using the fixed-effects model *∼ClinGene_DEG_group + Sex + Age_at_death + PMI_hrs*, with separate contrasts computed for sporadic and *C9orf72* FTD & FTD-MND relative to controls. Significant neighborhoods were defined at SpatialFDR < 0.1. Robustness analyses were additionally performed across multiple combinations of neighborhood size (k = 50, 100, 200, 300, 500, 750) and neighborhood sampling proportion (prop = 0.001, 0.01, 0.05, 0.1, 0.2).

We evaluated a range of neighborhood sizes and sampling proportions based on Milo recommendations and observed highly consistent results across parameter combinations and between LN10-enriched and unenriched populations (Extended Data Fig. 9a-e). Given this robustness, subsequent analyses used k = 150 and prop = 0.1 (Extended Data Fig. 9f-g). Differential abundance patterns were also consistent between grouped FTD/FTD-MND cases and analyses performed separately on FTD-only and FTD-MND-only cases (Extended Data Fig. 9h-i). In contrast, MND-only cases showed minimal FI depletion, consistent with the known regional vulnerability profile of MND (Extended Data Fig. 9j).

To identify disease- or control-specific effects on DA, we created a new metadata column combining the existing “batch” and “Individual” columns, and passed it to sample_col. In addition, we combined the sort category (Hoechst or LN10) with the Clinical Grouping x Gene Status information into a new column “sort_genotype”. We then specified design=∼sort_genotype and defined the contrasts depending on the question of interest - for example, to inspect differential abundance between C9 and control cases only across LN10-enriched cells we used sort_genotypeLN10_FTD__FTD_MND_C9-sort_genotypeLN10_Control.

### SCENIC analysis

To identify potential transcriptional regulators through the correlation of transcription factor (TF) expression with the expression of TF targets (regulons) we applied pySCENIC^71^ to the L5 ET subcluster. For the list of transcription factors we used the allTFs_hg38.txt file provided on the GitHub page of the Aerts Lab (https://github.com/aertslab). We ran the algorithm retaining only the 2000 most highly variable genes between the three L5 ET subclusters, in addition to any remaining TF not in the set of 2000. To perform the analysis we utilized the log transformed counts matrix. To identify regulons with different distributions across the three L5 ET subclusters, we applied the Kruskal-Wallis non-parametric test.

### Correlating differential abundance with neuronal size

To assess the relationship between neuronal size and selective vulnerability, neighborhood-level differential abundance estimates were correlated with neighborhood-level LN10 enrichment scores. Milo neighborhoods were generated in the scVI latent space using a k-nearest neighbor graph (n_neighbors = 150) and prop = 0.1. To estimate relative neuronal size via sort enrichment for different disease/genotype categories, differential abundance testing was first performed comparing LN10 and Hoechst conditions for control samples only using the contrast=genotype_sortControl_LN10 - genotype_sortControl_Hoechst, with neighborhood counts aggregated by individual-batch combination. Disease-associated differential abundance was subsequently computed separately for sporadic and *C9orf72* FTD & FTD-MND relative to controls following neighborhood annotation using the cell level 3 taxonomy. Pearson correlations were then computed between LN10-versus-Hoechst log fold-change values and disease-associated differential abundance log fold-change values across excitatory neuron neighborhoods and within individual excitatory neuron subclasses.

### Vulnerability-associated transcriptional programs

To identify baseline molecular programs associated with selective vulnerability, neighborhood-level depletion scores derived from Milo disease comparisons were regressed against average gene expression of control cells in the same neighborhoods. Briefly, Milo neighborhoods were generated as described above from the scVI latent space using a k-nearest neighbor graph (k = 150), and differential abundance statistics were computed separately for sporadic and *C9orf72* FTD & FTD-MND relative to controls while controlling for sex, age at death and PMI. Control excitatory neuron neighborhoods were then pseudobulked by averaging gene expression across constituent cells within each neighborhood. Pearson correlations were computed between baseline neighborhood-level gene expression and disease-associated neighborhood log fold-change depletion scores, across excitatory neuron neighborhoods. Genes whose baseline expression negatively correlated with depletion were interpreted as vulnerability-associated, whereas positively correlated genes were interpreted as resilience-associated. Gene set enrichment analyses and over-representation analyses were subsequently performed on vulnerability- and resilience-associated gene sets. For GSEA, genes were ranked by their correlation with depletion.

### RNAscope fluorescent in situ hybridization

Multiplex RNAscope fluorescent in situ hybridization was performed on human FI tissue sections using an RNAscope™ Multiplex Fluorescent Reagent Kit v2 (ACD Bio-Techne, catalog #323100) according to the manufacturer’s instructions. Candidate L5 ET subtype markers (*OPN4*: catalog #504991; *GADL1*: catalog #1617141; *TMPRSS15*: catalog # 545811; *RREB1*: catalog #570111) were evaluated together with *GABRQ* RNA labeling (Catalog # 483171-C3) and NEUN immunostaining. For each section, regions with *GABRQ*-positive neurons were first identified and, together with DAPI signal, used to identify cortical layer 5. Next, multiple separate z-stacked images were taken per slide capturing all channels in cortical layer V. Neurons were identified on the basis of NEUN signal. Morphologically defined von Economo neurons (VENs) were identified on the basis of NEUN and *GABRQ*-expression combined with established histological criteria. *GABRQ*+ neurons that were not morphologically VENs were defined as non-VEN *GABRQ*+. Neurons that did not express *GABRQ* were defined as neighboring neurons (NN). Marker expression frequencies were quantified across VENs, non-VEN *GABRQ*+ neurons, and neighboring neurons in a blinded manner such that the investigator that assessed signal of L5 ET marker genes per cell was agnostic of channels other than Hoechst, nor knew whether a cell was morphologically defined as a VEN.

### Statistical analysis

Statistical analyses were performed in R and Python. Unless otherwise specified, multiple-testing correction was performed using the Benjamini-Hochberg false discovery rate procedure. Pearson correlation coefficients were used to assess concordance between transcriptional signatures and differential abundance estimates. Exact statistical tests, sample sizes, and significance thresholds are provided in the corresponding figure legends and Supplementary Tables.

