## Supplementary material for "Single-cell transcriptomic atlas of frontoinsular cortex reveals molecular correlates of selective neuronal vulnerability in FTD": Suppl.Table_1: Table_1_Documentation.docx

*Written by: Arnar Breevoort*

*Last updated: April 27th, 2026*

**General:** This document explains the contents of Tables 1A, 1B and 1C.

Below is information regarding Table 1A

| **Column Name** | **Description** |
| --- | --- |
| Case ID | Anonymized ID for each case in the cohort |
| Gene status (C9orf72, sporadic) | Each disease case in the cohort has been categorized as either a *C9orf72*-mutation carrier or sporadic. Control cases don’t have a gene status category |
| Sex | Sex of each donor (M= male; F=female) |
| Age at Death | Age at death of donor (in years) |
| Clinical diagnosis | Clinical diagnosis of each donor |
| Primary neuropath diagnosis | Primary neuropathological diagnosis for each donor that was established postmortem |
| Grouping ClinDx | Broad clinical diagnosis grouping of each donor. Some donors have only FTD. Others have a spectrum of FTD-MND. Some have only MND. |
| ClinGene_DEG_group | Combined broad clinical diagnosis category (Grouping ClinDx) with Gene status. This category was used for downstream Dreamlet differential gene expression analysis |
| Grouping NPDx | Broad neuropathological grouping of each donor |
| PMI (hrs) | Post mortem interval between death of donor and preservation of brain samples |
| RIN | RNA integrity number (RIN) for frontoinsular cortex samples that were studied from each donor. RINs were established prior to the generation of single cell transcriptomics data and all passed quality control with the exception of Case ID 28 |
| First braincutting protocol | Braincutting protocol that was used to prepare brain samples |
| ADNC level | ADNC level of each donor |
| AD Thal Phase | AD Thal Phase of each donor |
| AD Braak Stage | AD Braak Stage of each donor |
| AD CERAD neuritic plaque score | AD CERAD neuritic plaque score of each donor |
| LBD stage (brainstem, transitional, diffuse) | LBD stage of each donor |
| neurodegen_score | Composite semiquantitative score of neurodegeneration severity in the ACC. Different features were assigned semiquantitative scores, and composite was calculated as: ((Vacuolation + Gliosis) / 2) + Neuronal Loss |
| TDP_score | Composite semiquantitative score of TDP-43 pathology in the ACC. Different features were assigned semiquantitative scores, and composite was calculated as: ((Dystrophic Neurites + Glial Cytoplasmic Inclusions + White Matter Dots or Threads) / 3) + Neuronal Cytoplasmic Inclusions |
| composite_score | Composite of neurodegeneration and TDP-43 severity scores above. Calculated as: neurodegen_score + TDP_score |
| CDR_Box_latest | Latest patient Sum of Boxes score on the standard Clinical Dementia Rating test. Scores range from 0-18, where higher score indicates greater impairment. |
| CDR_FTLD.Box_latest | Latest patient Sum of Boxes score on the modified Clinical Dementia Rating plus NACC FTLD test. Scores range from 0-24, where higher score indicates greater impairment. |
| MMSE_Total_latest | Latest patient score on the Mini-Mental State Examination. Ranges from 0-30, where scores below 23 indicate cognitive impairment. |

Below is information regarding Table 1B

| **Column** | **Description** |
| --- | --- |
| cell_level_1 | Name of cell type when using our most coarse taxonomy (level 1) |
| cell_level_2 | Name of cell type defined by reference mapping to Allen Brian Atlas 10x Whole Human Brain taxonomy and the 10x Human MTG SEA-AD taxonomy using MapMyCells (level 2) (Methods) |
| cell_level_3 | Name of cell type when using our most granular taxonomy (level 3) |
| cell_number | Total number of cells within each cell class |
| Sort Condition | Proportion of cells within each cell class that are from the non-enriched Hoechst sorting condition and from the enriched LN10 sorting condition. As expected, excitatory neurons are mostly derived from the LN10 sorting condition |
| Clinical Condition | Proportion of cells within each cell class that are from each Clinical Condition. ClinGene_DEG_group are used here as clinical conditions |
| Sex | Proportion of each cells within each cell class that are from female of male donors |
| Total counts median | Median total counts for all cells within each cell class |
| number of genes detected - median | Median number of genes that are detected per cell within each cell class |
| Percentage mitochondrial reads median | Median percentage of mitochondrial reads per cell within each cell class |

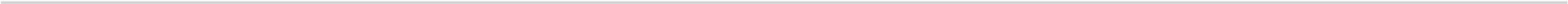

Below is information regarding Table 1C

| **Column** | **Description** |
| --- | --- |
| names | Names of genes for which logreg values are given. Not that the base script to generate this analysis can be found on our Github (Figure 1/run_rank_genes_groups_leiden2_061225.py) |
| scores | Logreg score for each gene listed in *names* |
| Leiden_2 | Cluster name that was in the original logreg script (Figure 1/run_rank_genes_groups_leiden2_061225.py). For clarity, we appended the final cell_level_3 classification names in column *cell_level_3* |
| cell_level_3 | Cell_level_3 classifications |

Below is information regarding Table 1D

| **Column** | **Description** |
| --- | --- |
| names | Names of genes for which Wilcoxon values are given. Not that the base script to generate this analysis can be found on our Github (Figure 1/run_rank_genes_groups_leiden2_061225.py) |
| scores | wilcoxon score for each gene listed in *names* for the cell type listed in column *cell_level_3* |
| logfoldchange | Relative expression value of gene listed in *names* for the cell type listed in *cell_level_3* |
| Leiden_2 | Cluster name that was in the original script (Figure 1/run_rank_genes_groups_leiden2_061225.py). For clarity, we appended the final cell_level_3 classification names in column *cell_level_3* |
| cell_level_3 | Cell_level_3 classifications |

Below is information regarding Table 1E

| **Column** | **Description** |
| --- | --- |
| names | Names of genes for which t-test values are given. Not that the base script to generate this analysis can be found on our Github (Figure 1/run_rank_genes_groups_leiden2_061225.py) |
| scores | t-test score for each gene listed in *names* for the cell type listed in column *cell_level_3* |
| logfoldchange | Relative expression value of gene listed in *names* for the cell type listed in *cell_level_3* |
| Leiden_2 | Cluster name that was in the original script (Figure 1/run_rank_genes_groups_leiden2_061225.py). For clarity, we appended the final cell_level_3 classification names in column *cell_level_3* |
| cell_level_3 | Cell_level_3 classifications |
