## Supplementary material for "Single-cell transcriptomic atlas of frontoinsular cortex reveals molecular correlates of selective neuronal vulnerability in FTD": Suppl.Table_3: Table_3_Documentation.docx

*Written by: Arnar Breevoort*

*Last updated: April 27th, 2026*

**General:** This document explains the contents of Suppl. Tables 3a and 3b. The scripts used to generate these tables can be found on our Github (see below).

**Scripts**

| **Table name** | **Script used to generate table** |
| --- | --- |
| Suppl. Table 3a | Milo Differential Abundance Analysis |
| Suppl. Table 3b | Gene Expression - Differential Abundance correlation analysis |

**Github:** <https://github.com/Pollen-lab/Transcriptional_Atlas_Frontoinsular_Cortex_in_ALS_and_FTD.git>.

Below is information regarding Suppl. Table 3a

| **Column Name** | **Description** |
| --- | --- |
| Neighborhood | Name of the Milo neighborhood. Milo generates neighborhoods of cells which are used for downstream analyses |
| C9_logFC | For each given neighborhood, there is a C9_logFC score which represents how differentially abundant that neighborhood is in C9orf72 FTD__FTD-MND compared to control. A negative value indicates depletion in disease. A positive value indicates enrichment in disease |
| S_logFC | For each given neighborhood, there is a S_logFC score which represents how differentially abundant that neighborhood is in sporadic FTD__FTD-MND compared to control. A negative value indicates depletion in disease. A positive value indicates enrichment in disease |
| sort_logFC | For each given neighborhood, there is a sort_logFC score which represents how differentially abundant that neighborhood is in the LN10 sorting condition compared to the Hoechst sorting condition. A negative value indicates that a neighborhood is depleted in the LN10 sort compared to the Hoechst sort. A positive value indicates that the neighborhood is enriched in the LN10 sort. Since the LN10 sort is gated on the basis of nuclear size, a positive sort_logFC score is a marker for large nuclei |
| C9_spatialFDR | C9_spatialFDR represents the spatial FDR value for the C9_logFC value for each neighborhood. Milo uses an adaptation of the Spatial FDR correction introduced by cydar, where p-values are corrected accounting for the amount of overlap between neighbourhoods. Specifically, each hypothesis test p-value is weighted by the reciprocal of the kth nearest neighbour distance. Milo recommends using spatial FDR over FDR. |
| S_spatialFDR | S_spatialFDR represents the spatial FDR value for the S_logFC value for each neighborhood |
| sort_spatialFDR | sort_spatialFDR represents the spatial FDR value for the sort_logFC value for each neighborhood |
| C9_FDR | C9_FDR represents the standard Benjamini-Hochberg correction FDR value for the C9_logFC value for each neighborhood. Milo recommends using spatial FDR instead. |
| S_FDR | S_FDR represents the standard Benjamini-Hochberg correction FDR value for the S_logFC value for each neighborhood. Milo recommends using spatial FDR instead. |
| sort_FDR | sort_FDR respresents the standard Benjamini-Hochberg correction FDR value for the sort_logFC value for each neighborhood. |
| nhood_annotation | For each neighborhood, we determine the dominant cell class measured by the majority of cells that are from a given cell class in each neighborhood. When nhood_annotation is L5_ET, this means that the majority of cells within that neighborhood have cell annotation “L5_ET” |

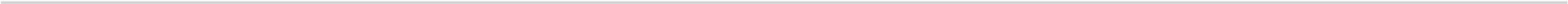

Below is information regarding Suppl. Table 3b

| **Column Name** | **Description** |
| --- | --- |
| gene_symbols | Gene names |
| Correlation | For each neighborhood, we calculated the correlation between gene expression in control cells with the differential abundance score of that neighborhood in disease. For each gene, there is a correlation value between gene expression (GE) and differential abundance (DA).  Note that a negative correlation value indicates that high expression of this gene in control cells correlates with a negative DA score (i.e. depletion) in disease and is thus an indication of vulnerability. A positive correlation indicates that high expression of this gene correlates with a positive DA score (i.e. enrichment) in disease and is thus an indication of resilience. |
| Correlation Gene Expression in control cells - Differential Abundance in C9orf72 FTD&FTD-MND | C9orf72 FTD&FTD-MND indicates that these calculations were done using Milo logFC DA scores from C9orf72 FTD&FTD-MND samples. |
| Correlation Gene Expression in control cells - Differential Abundance in Sporadic FTD&FTD-MND | Sporadic FTD&FTD-MND indicates that these calculations were done using Milo logFC DA scores from Sporadic FTD&FTD-MND samples. |
