## Supplementary material for "Single-cell transcriptomic atlas of frontoinsular cortex reveals molecular correlates of selective neuronal vulnerability in FTD": Suppl.Table_4: Table_4_Documentation.docx

*Written by: Arnar Breevoort*

*Last updated: April 27th, 2026*

**General:** This document explains the contents of Suppl. Tables 3a and 3b. The scripts used to generate these tables can be found on our Github (see below).

**Scripts**

| **Table name** | **Script used to generate table** |
| --- | --- |
| Suppl. Table 4a-c | Figure 4/Cell_Marker_Detection_L5ET_Subtypes.ipynb |

**Github:** <https://github.com/Pollen-lab/Transcriptional_Atlas_Frontoinsular_Cortex_in_ALS_and_FTD.git>.

Below is information regarding Table 4a

| **Column** | **Description** |
| --- | --- |
| names | Names of genes for which logreg values are given. Not that the base script to generate this analysis can be found on our Github (Figure 4/Cell_Marker_Detection_L5ET_Subtypes.ipynb) |
| scores | Logreg score for each gene listed in *names* |
| Cluster | Name of the L5 ET subtype. Note that classifications are already embedded in the column for clarification. |

Below is information regarding Table 4b

| **Column** | **Description** |
| --- | --- |
| names | Names of genes for which Wilcoxon values are given. Not that the base script to generate this analysis can be found on our Github (Figure 4/Cell_Marker_Detection_L5ET_Subtypes.ipynb) |
| scores | wilcoxon score for each gene listed in *names* for the cell type listed in column *cluster* |
| logfoldchange | Relative expression value of gene listed in *names* for the cell type listed in *cluster* |
| Cluster | Name of the L5 ET subtype. Note that classifications are already embedded in the column for clarification. |

Below is information regarding Table 4c

| **Column** | **Description** |
| --- | --- |
| names | Names of genes for which t-test values are given. Not that the base script to generate this analysis can be found on our Github (Figure 4/Cell_Marker_Detection_L5ET_Subtypes.ipynb) |
| scores | t-test score for each gene listed in *names* for the cell type listed in column *cluster* |
| logfoldchange | Relative expression value of gene listed in *names* for the cell type listed in *cell_level_3* |
| Cluster | Name of the L5 ET subtype. Note that classifications are already embedded in the column for clarification. |

Below is information regarding Table 4d

| **Column** | **Description** |
| --- | --- |
| **Note that this table contains the raw counts that were determined for the L5ET RNAscope analysis. Quantifications were done in a blinded manner (see methods)** | |
| Image No. | Multiple images were taken from each sample used to examine L5ET markers. The number here represents the image number for each of the examined L5ET markers. |
| Neuron No | For each image, neurons were quantified on the basis of NEUN signal (protein). The number represents neurons for each image |
| VEN | Von Economo neurons (VENs) were identified on the basis of NEUN signal (protein), GABRQ signal (RNAscope) and morphology (see methods) |
| Fork cell | Fork cells were identified on the basis of NEUN signal (protein), and morphology (see methods) |
| NN | Neighboring neurons were defined as neurons that were not VENs or Fork cells |
| Unclassifiable | Unclassifiable were neurons that had characteristics of VENs or fork cells, but for which we could not determine identity with confidence. Generally, this was due to the cell body of the neuron not being fully capture in the imaging z-stack, limiting confidence in the complete morphology of the neuron. Oftentimes, these very well could have been VENs |
| GABRQ positive | Neurons with positive GABRQ (RNAscope) signal |
| L5ET Marker (e.g. OPN4) | Neurons with positive L5ET marker signal (e.g. OPN4) |

Below is information regarding Table 4e

| **Column** | **Description** |
| --- | --- |
| **Note that this table contains summarized counts of table 4d that were used to plot the barplots as seen in Figure 4 and Extended data fig. 14** | |
| Marker | L5ET marker that was summarized in the sub table |
| Category | Category of cell type as defined in the raw counts table. To determine the quality of the L5ET marker, we focussed on three main categories. These were VENs that express GABRQ, non-VENs that express GABRQ, and Neighboring Neurons that do not express GABRQ. A proper VEN marker would differentiate VENs from non-VEN GABRQ cells |
| Image index | Image number as described in the raw counts table |
| Pos | Number of positive cells for each category within the given image index. Note that for some images there were multiple VENs or other cell types |
| Total | Number of cells total for each category within the given image index. Note that for some images there were multiple VENs or other cell types |
| PCT | Percentage of all cells for each category within given image index that were positive for the L5ET marker as described in column *marker* |
