## Supplementary material for "Single-cell transcriptomic atlas of frontoinsular cortex reveals molecular correlates of selective neuronal vulnerability in FTD": Suppl_Table_2_Documentation

*Written by: Arnar Breevoort*

*Last updated: April 27th, 2026*

**General:** This document explains the contents of the differential gene expression (DGE) matrices that are part of table 2. The DGE data is generated using Dreamlet (1) (see methods and scripts).

**Note:** since the DGE files exceed the maximum file size for a BioRxiv submission, the files that are part of Suppl. Table 2 can be accessed here: <https://ucsf.box.com/s/04230hrm0r9f6wqnuekgapm1q5k5iutc>

Below is information regarding the different csv files with DGE data from Suppl. Tables 2a-b

| **File Name** | **Description** |
| --- | --- |
| dreamlet_DEG_ALL_C9-FTD_FTD-MND_v_Control_LV1 | DGE comparison between **FTD__FTD_MND_C9** cases and **Control** cases using taxonomy level cell level 1 |
| dreamlet_DEG_ALL_C9-FTD_FTD-MND_v_Control_LV2 | DGE comparison between **FTD__FTD_MND_C9** cases and **Control** cases using taxonomy level cell level 2 |
| dreamlet_DEG_ALL_C9-FTD_FTD-MND_v_Control_LV3 | DGE comparison between **FTD__FTD_MND_C9** cases and **Control** cases using taxonomy level cell level 3 |
| dreamlet_DEG_ALL_Sporadic_MND_vs_Control_LV1 | DGE comparison between **FTD__FTD_MND_sporadic** cases and **Control** cases using taxonomy level cell level 1 |
| dreamlet_DEG_ALL_Sporadic_MND_vs_Control_LV2 | DGE comparison between **FTD__FTD_MND_sporadic** cases and **Control** cases using taxonomy level cell level 2 |
| dreamlet_DEG_ALL_Sporadic_MND_vs_Control_LV3 | DGE comparison between **FTD__FTD_MND_sporadic** cases and **Control** cases using taxonomy level cell level 3 |
| dreamlet_DEG_ALL_C9_MND_vs_Control_LV1 | DGE comparison between **MND_C9** cases and **Control** cases using taxonomy level cell level 1 |
| dreamlet_DEG_ALL_C9_MND_vs_Control_LV2 | DGE comparison between **MND_C9** cases and **Control** cases using taxonomy level cell level 2 |
| dreamlet_DEG_ALL_C9_MND_vs_Control_LV3 | DGE comparison between **MND_C9** cases and **Control** cases using taxonomy level cell level 3 |
| dreamlet_DEG_ALL_Sporadic_MND_vs_Control_LV1 | DGE comparison between **MND_sporadic** cases and **Control** cases using taxonomy level cell level 1 |
| dreamlet_DEG_ALL_Sporadic_MND_vs_Control_LV2 | DGE comparison between **MND_sporadic** cases and **Control** cases using taxonomy level cell level 2 |
| dreamlet_DEG_ALL_Sporadic_MND_vs_Control_LV3 | DGE comparison between **MND_sporadic** cases and **Control** cases using taxonomy level cell level 3 |
| dreamlet_DEG_ALL_S_FTD_FTD-MND_vs_C9_FTD_FTD-MND_LV1 | DGE comparison between **MND_sporadic** cases and **MND_C9** cases using taxonomy level cell level 1 |
| dreamlet_DEG_ALL_S_FTD_FTD-MND_vs_C9_FTD_FTD-MND_LV2 | DGE comparison between **MND_sporadic** cases and **MND_C9** cases using taxonomy level cell level 2 |
| dreamlet_DEG_ALL_S_FTD_FTD-MND_vs_C9_FTD_FTD-MND_LV3 | DGE comparison between **MND_sporadic** cases and **MND_C9** cases using taxonomy level cell level 3 |
| dreamlet_DEG_ALL_Sporadic-FTD_FTD-MND_v_Control_LV1 | DGE comparison between **FTD__FTD_MND_sporadic** cases and **FTD__FTD_MND_C9** cases using taxonomy level cell level 1 |
| dreamlet_DEG_ALL_Sporadic-FTD_FTD-MND_v_Control_LV2 | DGE comparison between **FTD__FTD_MND_sporadic** cases and **FTD__FTD_MND_C9** cases using taxonomy level cell level 2 |
| dreamlet_DEG_ALL_Sporadic-FTD_FTD-MND_v_Control_LV3 | DGE comparison between **FTD__FTD_MND_sporadic** cases and **FTD__FTD_MND_C9** cases using taxonomy level cell level 3 |

Below is information regarding relevant columns in the DGE matrices from Suppl. Tables 2a-b

| **Column** | **Description** |
| --- | --- |
| **logFC** | LogFC of given gene in disease compared to control. (log2-transformed fold changes) |
| **P.Value** | P.Value |
| **adj.P.Val** | Adjusted.P.Value |
| **assay** | Cell type (Ex = excitatory neuron; In = inhibitory neuron; NN=non-neuronal) |
| **genes** | Gene for which DGE output is given in that row |
| **coef** | Groups compared (FTD__FTD_MND_C9_vs_Control= C9orf72 FTD & FTD-MND compared to control; FTD__FTD_MND_sp_vs_Control = sporadic FTD & FTD-MND compared to control; FTD_S_vs_FTD_C9 = sporadic FTD & FTD-MND compared to C9orf72 FTD & FTD-MND |

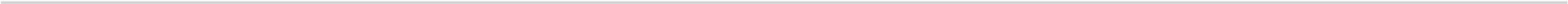

Below is information regarding the different csv files with Zenith data from Suppl. Tables 2c

| **File Name** | **Description** |
| --- | --- |
| FTD_&FTD-MND_C9_vs_C_level_1_res_zenith_BP | Zenith results from comparisons between **FTD__FTD_MND_C9** cases and **Control** cases using taxonomy level cell level 1 [Biological Processes] |
| FTD_&FTD-MND_C9_vs_C_level_1_res_zenith_CC | Zenith results from comparisons between **FTD__FTD_MND_C9** cases and **Control** cases using taxonomy level cell level 1 [Cellular Components] |
| FTD_&FTD-MND_C9_vs_C_level_1_res_zenith_MF | Zenith results from comparisons between **FTD__FTD_MND_C9** cases and **Control** cases using taxonomy level cell level 1 [Molecular Function] |
| FTD_&FTD-MND_C9_vs_C_level_2_res_zenith_CC | Zenith results from comparisons between **FTD__FTD_MND_C9** cases and **Control** cases using taxonomy level cell level 2 [Cellular Components] |
| FTD_&FTD-MND_C9_vs_C_level_2_res_zenith_MF | Zenith results from comparisons between **FTD__FTD_MND_C9** cases and **Control** cases using taxonomy level cell level 2 [Molecular Function] |
| FTD_&FTD-MND_C9_vs_C_level_3_res_zenith_BP | Zenith results from comparisons between **FTD__FTD_MND_C9** cases and **Control** cases using taxonomy level cell level 3 [Biological Processes] |
| FTD_&FTD-MND_C9_vs_C_level_3_res_zenith_CC | Zenith results from comparisons between **FTD__FTD_MND_C9** cases and **Control** cases using taxonomy level cell level 3 [Cellular Components] |
| FTD_&FTD-MND_C9_vs_C_level_3_res_zenith_MF | Zenith results from comparisons between **FTD__FTD_MND_C9** cases and **Control** cases using taxonomy level cell level 3 [Molecular Function] |
| FTD_&FTD-MND_S_vs_C_level_1_res_zenith_BP | DGE comparison between **FTD__FTD_MND_sporadic** cases and **Control** cases using taxonomy level cell level 1 [Biological Processes] |
| FTD_&FTD-MND_S_vs_C_level_1_res_zenith_CC | DGE comparison between **FTD__FTD_MND_sporadic** cases and **Control** cases using taxonomy level cell level 1 [Cellular Components] |
| FTD_&FTD-MND_S_vs_C_level_1_res_zenith_MF | DGE comparison between **FTD__FTD_MND_sporadic** cases and **Control** cases using taxonomy level cell level 1 [Molecular Function] |
| FTD_&FTD-MND_S_vs_C_level_2_res_zenith_BP | DGE comparison between **FTD__FTD_MND_sporadic** cases and **Control** cases using taxonomy level cell level 2 [Biological Processes] |
| FTD_&FTD-MND_S_vs_C_level_2_res_zenith_CC | DGE comparison between **FTD__FTD_MND_sporadic** cases and **Control** cases using taxonomy level cell level 2 [Cellular Components] |
| FTD_&FTD-MND_S_vs_C_level_2_res_zenith_MF | DGE comparison between **FTD__FTD_MND_sporadic** cases and **Control** cases using taxonomy level cell level 2 [Molecular Function] |
| FTD_&FTD-MND_S_vs_C_level_3_res_zenith_BP | DGE comparison between **FTD__FTD_MND_sporadic** cases and **Control** cases using taxonomy level cell level 3 [Biological Processes] |
| FTD_&FTD-MND_S_vs_C_level_3_res_zenith_CC | DGE comparison between **FTD__FTD_MND_sporadic** cases and **Control** cases using taxonomy level cell level 3 [Cellular Components] |
| FTD_&FTD-MND_S_vs_C_level_3_res_zenith_MF | DGE comparison between **FTD__FTD_MND_sporadic** cases and **Control** cases using taxonomy level cell level 3 [Molecular Function] |

Below is information regarding the different csv files with Zenith data from Suppl. Tables 2c

| **Column** | **Description** |
| --- | --- |
| **assay** | Cell type from cell level 1, cell level 2 or cell level 3 taxonomies (see Suppl. Table 1b) |
| **coef** | Zenith results are from comparisons made between the listed coefficients. Note that FTD_S_vs_C represents a comparison between the sporadic FTD & FTD-MND clinical category with control. FTD_C9_vs_C represents a comparison between the *C9orf72* FTD & FTD-MND clinical category with control. |
| **Geneset** | GO terms |
| **Delta** | **Estimated average directional effect** of the genes in a gene set relative to all other genes in the analysis |
| **NGenes** | Number of genes within each GO term listed in the Geneset column |
| **Direction** | GO term can either by up or down in disease compared to control |
| **FDR** | FDR value for shown delta values |

**References**

(1) <https://diseaseneurogenomics.github.io/dreamlet/>
